# Perivascular Macrophages Convert Physical Wound Signals Into Rapid Vascular Responses

**DOI:** 10.1101/2024.12.09.627538

**Authors:** Zaza Gelashvili, Zhouyang Shen, Yanan Ma, Mark Jelcic, Philipp Niethammer

## Abstract

Leukocytes detect distant wounds within seconds to minutes, which is essential for effective pathogen defense, tissue healing, and regeneration. Blood vessels must detect distant wounds just as rapidly to initiate local leukocyte extravasation, but the mechanism behind this immediate vascular response remains unclear. Using high-speed imaging of live zebrafish larvae, we investigated how blood vessels achieve rapid wound detection. We monitored two hallmark vascular responses: vessel dilation and serum exudation. Our experiments—including genetic, pharmacologic, and osmotic perturbations, along with chemogenetic leukocyte depletion—revealed that the cPla_2_ nuclear shape sensing pathway in perivascular macrophages converts a fast (∼50 μm/s) osmotic wound signal into a vessel-permeabilizing, 5-lipoxygenase (Alox5a) derived lipid within seconds of injury.

These findings demonstrate that perivascular macrophages act as physicochemical relays, bridging osmotic wound signals and vascular responses. By uncovering this novel type of communication, we provide new insights into the coordination of immune and vascular responses to injury.

## Introduction

Realtime imaging of zebrafish larvae allows studying the spatiotemporal regulation of wound detection *in vivo* at timescales inaccessible to conventional biochemical or molecular biology methods^1, 2^. Previous work in zebrafish, flies, and frogs, revealed extracellular nucleotides, bioactive lipids, Ca^2+^ waves, reactive oxygen species-(ROS), osmotic-, and ionicgradients as physiological mediators of rapid wound detection^3–12^.

We previously showed that zebrafish detect integumental breaches via the osmotic shock that occurs when environmental freshwater enters their tissues from the outside. Consequently, immersing zebrafish larvae in isotonic salt or sugar solutions suppresses neutrophil recruitment and acute breach closure by epithelial cells. Compared to other isotonic treatments, NaCl has a stronger inhibitory effect^4, 5^, which may be due to electric fields (EFs)^7^, or other, yet unknown salt-sensing mechanisms. Osmotic shock at the wound induces local cell swelling and nuclear deformation that stretches the inner nuclear membrane (INM)^13^. Along with Ca^2+^, INM tension (T_INM_) recruits cPla_2_ to the INM to release arachidonic acid (AA). Lipoxygenases, cyclooxygenases, etc., further oxidize AA into bioactive lipids that control chemotaxis, cell differentiation, blood vessel tone and permeability, and many other processes^14–18^. Besides wound detection, the cPla_2_ nuclear shape sensing pathway also controls confined cell migration and the cell cycle^19–23^. At larval wounds and infection sites, downstream metabolites of the cPla_2._ nuclear shape sensing pathway rapidly recruits leukocytes^4, 24, 25^.

At least in part, these leukocytes come from the blood circuit. To release them near to wound sites, blood vessels must be able to sense wounds as fast or even faster than leukocytes. Although a plethora of vascular regulators have been described, the mechanisms that rapidly detect and relay tissue injury cues to nearby vessels remain largely unknown. Here, we report that perivascular macrophage nuclei mediate osmotic surveillance by blood vessels through the cPla_2_ nuclear shape sensing pathway.

## Results

### Osmotic surveillance mediates rapid wound detection by blood vessels

To image rapid vessel responses to injury *in vivo*, we intracardially injected 2-3 days post fertilization (dpf) larvae with 70 kDa fluorescent dextran to label the circulating blood pool. Carefully avoiding damage to the vasculature, we amputated the tail fin tips of these larvae in E3 solution^26^ adjusted to isotonicity with sodium chloride (ISO_NaCl_) to roughly match the composition of vertebrate interstitial fluid (**Fig. 1a**). We previously showed that ISO_NaCl_ prevents rapid leukocyte recruitment and epithelial wound closure in zebrafish larvae^4, 5, 27^. Wound detection can be initialized by transferring the injured larvae back to fresh water such as E3 (HYPO). This ISO→HYPO switching of bathing solutions at T= 270-300 s after starting the experiment (shift), temporally synchronizes the wound response in animals injured and imaged at different times, facilitating statistical integration. The individual contributions of osmotic pressure and ionic cues to wound detection can be distinguished by shifting into isotonic solutions of different salts or sugar compositions (ISO*).

**Figure 1.**
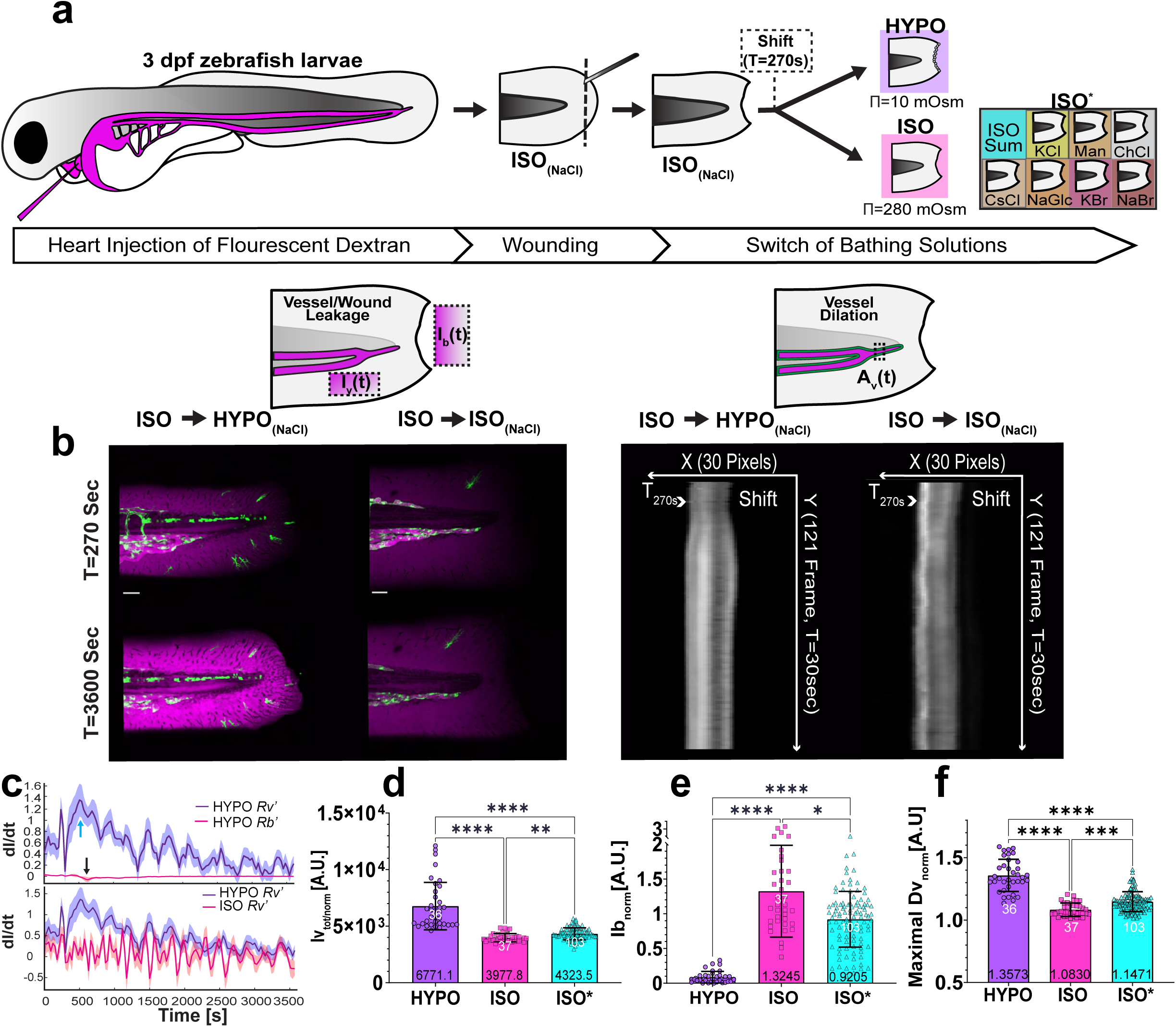
Osmotic surveillance mediates rapid wound detection by blood vessels. (**a**) Upper panel, cartoon scheme representing the experimental workflow of fluorescence microangiography, tailfin wounding, and ionic/osmotic bath treatment. HYPO, regular E3 solution. ISO_(NaCl)_, E3 adjusted to interstitial osmolarity by supplementing E3 with 135 mM NaCl. ISO*, E3 adjusted to isotonicity with other salts/osmolytes. Dotted box indicates time of solution switch (shift). (**b**) Cartoon schemes depicting the regions of interest for measuring the dextran permeability of vessels (Iv), wounds (Ib), as well as vessel dilation (Dv). Left panel, confocal maximal intensity projection (MIP) of wounded endothelial reporter larvae (Tg(*kdrl*:eGFP)) before (T= 270 s) and after (T= 3600 s) switch of bathing solutions. Green, *kdrl*:eGFP fluorescence. Magenta, pseudo-coloured 70 kDa dextran fluorescence. Right panel, kymographs of vessel diameter before and after switch of bathing solution. (**c**) Top panel, rate (dI dt^−1^) of vessel leakage (blue) and wound leakage (red) versus time upon switching from ISO_(NaCl)_ to HYPO. Arrows indicate the maximal rate of change for vessel permeability (cyan) or wound permeability (black). Bottom panel, rate (dI dt^−1^) of vessel leakage versus time upon switching from ISO_(NaCl)_ to HYPO (blue) or ISO_(NaCl)_ to ISO_(NaCl)_. Shaded error bars, SD. Note, the blue curves represent the same, replotted dataset. (**d**) Quantification of normalized, integrated (between T= 0-3600 s) dextran leakage (Iv_tot/norm_) or (**e**) or normalized dextran leakage from the wound (Ib_norm_) measured at T= 3600 s. (**f**) Maximal, normalized endothelial diameter (max(Dv_norm_)), obtained from kymograph). If indicated, the raw data was normalized by the mean of the first 10 frames (T= 0-270 s, i.e., preceding solution switching). Note, ISO* contains pooled results from different isotonic salt/osmolyte treatments. Error bars, SD. White numbers, animals. Black numbers, mean of dataset. P values, one-way analysis of variance (ANOVA) with Dunn’s post-hoc test (in **d, f, e**). All P values are listed in the Supplementary Excel File 1 (F1_T1). *P ≤ 0.05, **P ≤ 0.01, ***P ≤ 0.001, ****P ≤ 0.0001. Scale bars, 50 μm.

Using spinning disk confocal microscopy (**Fig. 1b left**), we followed vessel permeability by measuring the fluorescence intensity of 70 kDa fluorescent dextran in a region of interest (ROI) adjacent to the caudal vein (*Iv*(*t*)) (**Fig. 1b, top left cartoon**) after ISO-_NaCl_ → HYPO, ISO_NaCl_ → ISO_NaCl_, or ISO_NaCl_ → ISO* shifts. The apparent vessel leakage rate 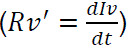 steeply rose immediately after switching from ISO_NaCl_ (T=0-270s) to HYPO bathing solution. Once dextran (as a proxy for serum) is exudated into the interstitial space, it may leak from the tissue into the bathing solution. To capture this “bleed out”, we monitored dextran intensity within a ROI on the external side of the amputation wound (*Ib*(*t*)) (**Fig. 1b, top left cartoon**). Unlike ISO_NaCl_ → ISO_NaCl_, ISO_NaCl_ → HYPO switching also induced rapid vessel dilation as measured by normalized vessel diameter increase(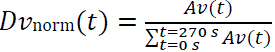) within a ROI comprising a vein/artery subsection (**Fig. 1b right, 1b top right cartoon**). The apparent wound leakage (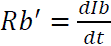) was negligible (*Rv*^′^ ≫ **R*b*′) compared to *Rv*^′^ (**Fig. 1c top panel)**. Furthermore, the *Rv*^′^ rate peaked at ∼500 s after the start of imaging, while *Rb*′ dropped with ∼2 min delay at ∼700 s. By contrast, replacing ISO_NaCl_ by fresh ISO_NaCl_ solution, did not cause vessel leakage of 70 kDa dextran compared to HYPO (**Fig. 1c bottom panel).** Apparent wound closure is inversely correlated to normalized wound permeability (calculated as 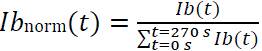) and was strongly inhibited by ISO_NaCl_, in line with previous data^5^. For systematic statistical comparison, we henceforth represent vessel leakage by 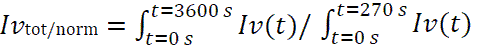 (**Fig. 1d, S1a**), overall wound closure as *Ib*_norm_(3600*s*) (**Fig.1e, S1b and Supplementary Video 1, 2**), and vessel dilation as max (*Dv*_norm_(*t*)), respectively (**Fig.1f, S1c**).

As observed earlier^4, 5^, ISO_NaCl_ showed a moderately stronger inhibition of some wound responses than some other ISO* treatments (**Fig.1e, S1d-f Supplementary Video 3**). At least in part, these differences might be explained by wound EF disruption as previously proposed^7^. The immediate increase of vessel permeability after injury points to a fast-propagating wound cue, such as osmotic shock resulting from NaCl leakage from the wound.

### Osmotic control of vessel permeability requires Alox5a

We previously showed that nuclear shape sensing by cPla_2_ mediates rapid osmotic wound detection by leukocytes via Alox5a^4, 6, 28, 29^. To test whether the cPla_2_-lipoxygenase pathway also mediates wound detection by blood vessels, we measured normalized, total vessel leakage (*Iv*_tot/norm_), and wound permeability (*Ib*_norm_(3600*s*)) upon genetic and pharmacological lipoxygenase perturbation. Neither parameter was significantly affected in *alox12*^mk215/mk215^ mutants as compared to wt animals. By contrast, *alox5a*^mk211/mk211^ and *lta_4_h*^mk216/mk216^ mutant larvae exhibited a significantly reduced vessel leakage upon ISO-_NaCl_ → HYPO shifting, whereas wound permeability was unaffected (**Fig. 2a-g and S2a-f**). Licofelone, a cyclooxygenase/5-lipoxygenase inhibitor^30^, and MK886, an inhibitor of 5-lipoxygenase activating protein (ALOX5AP)^30^, faithfully reproduced these genetic effects (**Fig. 2e-h and S2**). By contrast, cyclooxygenase inhibition with diclofenac did not inhibit vessel leakage.

**Figure 2.**
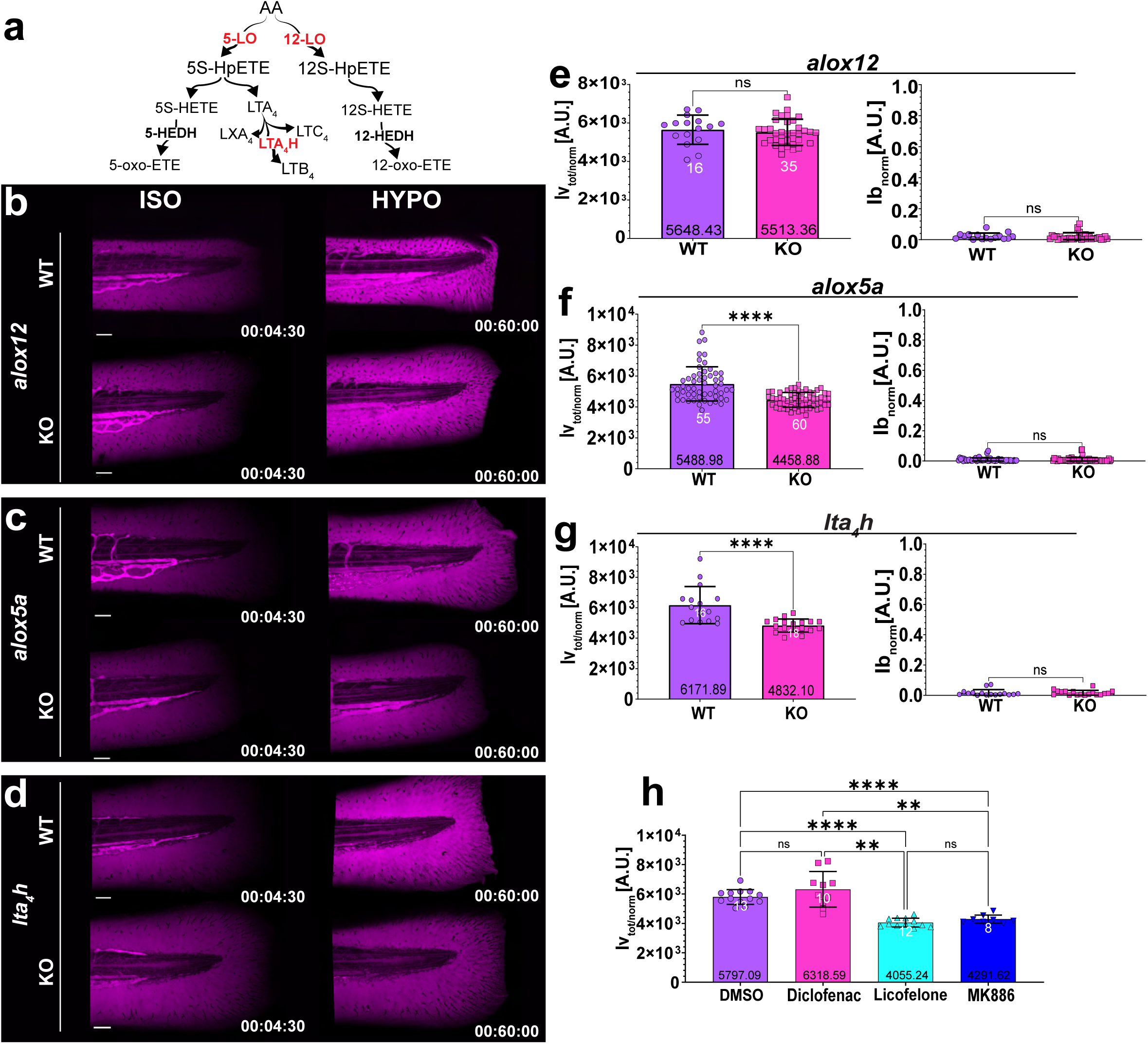
Osmotic control of vessel permeability requires Alox5a. (**a**) simplified scheme of enzymatic derivatives of AA, the pathways enzymes tested in the study are highlighted in red. Representative confocal MIP images of pseudo-coloured 70 kDa dextran fluorescence before (T=270 s, left panel) and after (T-3600 s, right panel) ISO_(NaCl)_ ◊ HYPO shift in wild-type animals (Casper), (**b**) *alox12*^mk215/mk215^ mutants, (**c**) *alox5a*^mk211/mk211^ mutants, (**d**) and *lta_4_h*^mk216/mk216^ mutants. 3 dpf larvae were compared to their matched wild-type siblings. (**e-g**) Quantification of integrated (T= 0-3600 s), normalized vessel leakage and wound permeability (T= 3600 s). The P values in (**e**) and (**g**) were calculated using a non-paired, two-tailed student’s t-test. The data in (**f**) were not normally distributed, and a two-sided Mann–Whitney U test was used instead. (**h**) Integrated vessel leakage after pretreatment of wild-type (Casper) larvae with 130 nM Diclofenac, 50 μM Licofelone and 10 μM MK886, or DMSO (vehicle) and ISO_(NaCl)_ ◊ HYPO shifting. P values, one-way ANOVA with Dunn’s post-hoc test. White numbers, animals. Black numbers, mean of dataset. Error bars, SD. All figure source data and numerical P values are listed in the Supplementary Excel File 1. **P ≤ 0.01, ****P ≤ 0.0001. ns, not significant (P > 0.05). Timestamp, hh: mm: ss. Scale bars, 50 μm.

Consistent with a role of the lipoxygenase pathway, bath supplementation with the cPla_2_ product AA partially (∼23%) restored vessel leakage in ISO_NaCl_ compared to hypotonic E3 media, without considerably altering wound closure or vessel dilation (**Fig. S3a-c**). Adenosine triphosphate (ATP), which we previously found to promote epithelial wound closure^5^, strongly blocked dextran leakage from wounds as expected (**Fig. S3b, e**). Yet, it only had a mild effect on vessel leakage and dilation. By contrast, adenosine affected dilation, but not vessel leakage and wound closure (**Fig. S3**). Although all these tissue damage responses are osmotically controlled, their underlying molecular mechanisms are different. cPla_2_ generates the AA for Alox5a, and directly translates osmotic wound cues into bioactive lipid signals^4, 6, 31^. Collectively, our data raised the possibility that nuclear membrane mechanotransduction via cPla_2_ contributed to rapid osmotic vessel permeabilization upon injury, but not to vessel dilation or wound closure. To further test this idea, we set out to identify the cells that were relaying the osmotic wound signal to the vessels.

### Macrophages mediate rapid osmotic vessel permeabilization

According to zebrafish single-cell RNA sequencing resources^32–34^, *alox5a* and *lta_4_h* are expressed in immune and non-immune (e.g., epithelial) cells. To distinguish whether vessel permeabilization through the Alox5a-Lta_4_h pathway is mediated by immune-or non-immune cells, we depleted neutrophils and macrophages, i.e., the two major classes of immune cells present in larval zebrafish tail fins, using the nitroreductase (NTR)/met-ronidazole cell ablation system (see methods)^35^. Neutrophil ablation did not modulate vessel permeability. So, the process of neutrophil extravasation (e.g., by poking holes into the vessel wall during passage) unlikely was responsible for the wound-dependent vessel leakage (**Fig. 3a-c**, **S4a, d; Supplementary Video 4**). Conversely, macrophage depletion significantly reduced vessel leakage (**Fig. 3b, d**, **S4b, d****; Supplementary Video 5**) upon ISO_NaCl_ → HYPO shifting just as global Alox5a-Lta_4_h pathway abrogation. Licofelone did not further block vessel permeabilization in macrophage-depleted animals, suggesting that macrophage Alox5a regulates acute vessel leakage (**Fig. 3e and S4c, f**). In line with previous data^36^, macrophage depletion did not affect acute neutrophil recruitment (**Fig. S4g, h**). Although both vessel permeabilization and innate immune defense in zebrafish larvae are governed by osmotic surveillance, they are carried out by distinct branches of the ALOX5 pathway. We and others previously showed that the acute neutrophil response to tissue injury and infection (i.e., within ∼ 2 h of injury) is independent of LTB4^24, 37^, and instead mediated by the alternative ALOX5 pathway product 5-KETE^4, 24^.

**Figure 3.**
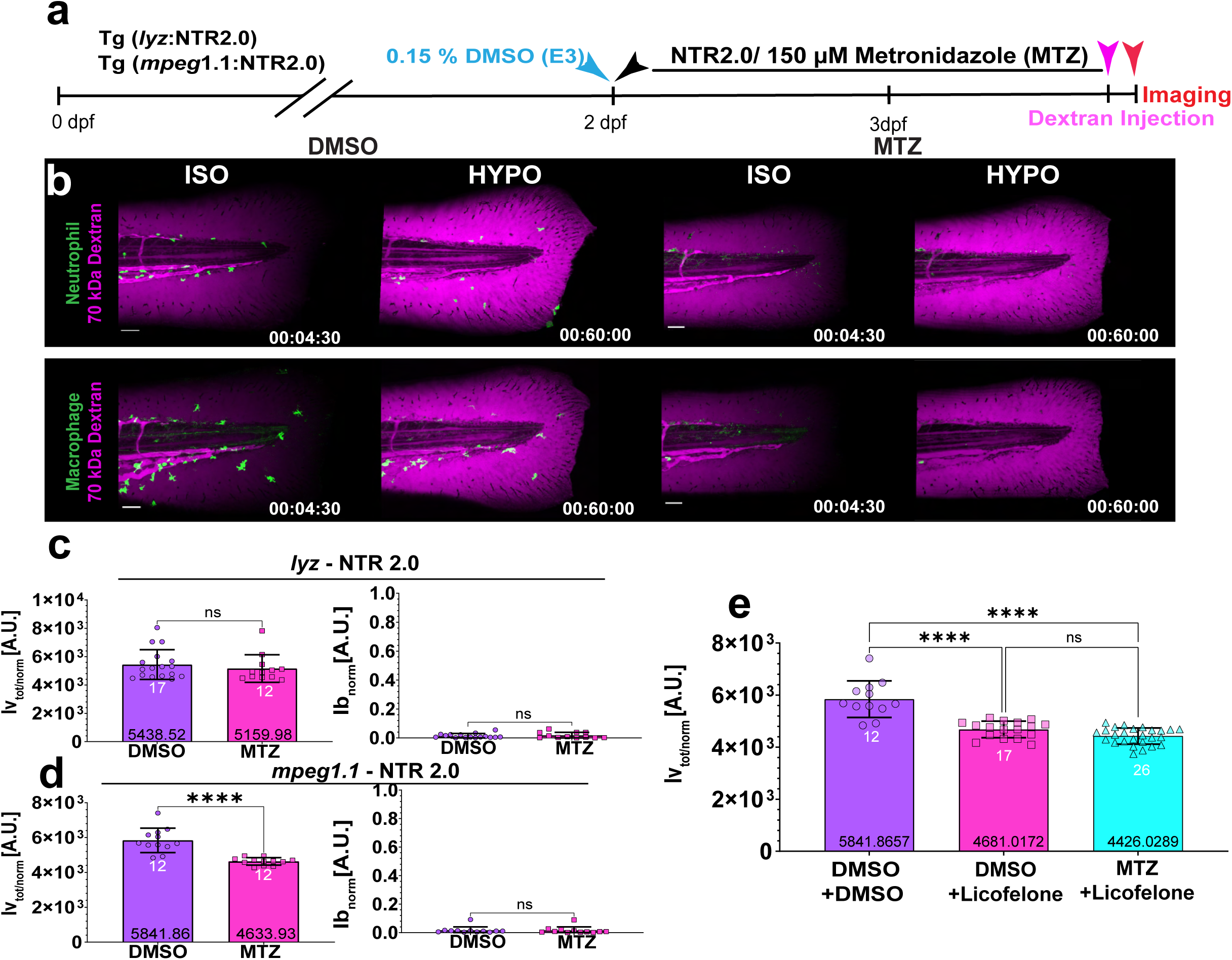
Macrophages mediate rapid osmotic vessel permeabilization. (**a**) Experimental timeline for metronidazole-induced depletion of macrophages or neutrophils in 3 dpf Tg(*mpeg1.1*: YFP-NTR2.0) or Tg(*lyz*: YFP-NTR2.0) larvae, respectively. Blue arrow, start of the vehicle treatment (0.15% DMSO). Black arrow, start of the metronidazole (150 μM MTZ) treatment. Magenta arrow, dextran injection. Red arrow, start of imaging. (**b**) Representative confocal MIPs of leukocytes and tissue dextran fluorescence before (T= 270 s) and after (T= 3600 s) ISO_(NaCl)_ → HYPO shifting. Green, pseudo-coloured *lyz:* YFP-NTR2.0 (upper panel) or *mpeg1.1*: YFP-NTR2.0 (lower panel) fluorescence. Magenta, pseudo-coloured 70 kDa dextran fluorescence. (**c**, **d**) Quantification of vessel and wound leakage in neutrophil-or macrophage-depleted animals. P values, non-paired, two-tailed Student’s t-test. (**e**) Quantification of vessel leakage after ISO_(NaCl)_ → HYPO shifting in macrophage-containing (DMSO + DMSO, DMSO + Licofelone) or –depleted (MTZ + Licofelone) larvae ± 50 μM Licofelone or 150 μM MTZ, respectively. P value, one-way ANOVA with Dunn’s post-hoc test. White numbers, animals. Black numbers, mean of dataset. Error bars, SD. The figure source data and numerical P values are listed in the Supplementary Excel File 1. ****P ≤ 0.0001. ns, not significant (P > 0.05) (**c-e**). Timestamp, hh:mm:ss. Scale bars, 50 μm.

### Nuclear shape sensing by cPla_2_ relays rapid osmotic wound cues to vessels

To test cPla_2_’s role in vessel permeabilization, we generated cPla_2_ mutant animals using CRISPR/Cas9 (**Fig.S5a, b**). Like the *alox5a* and *lta_4_h* mutants, cPla_2_ deficient larvae showed reduced vessel-but unaltered wound leakage upon ISO_NaCl_ → HYPO shifting (**Fig. 4a, S5c, d**). To directly test for cPla_2_ activation in macrophages, we expressed a fluorescently tagged version of the enzyme (cPla_2_-mKate2) using the *mpeg1.1* promoter. cPla_2_ activation is indicated by the enzymes’ rapid, mechanosensitive translocation from the nucleoplasm to the INM as we and others previously showed^4, 6, 20, 21, 31^.

**Figure 4.**
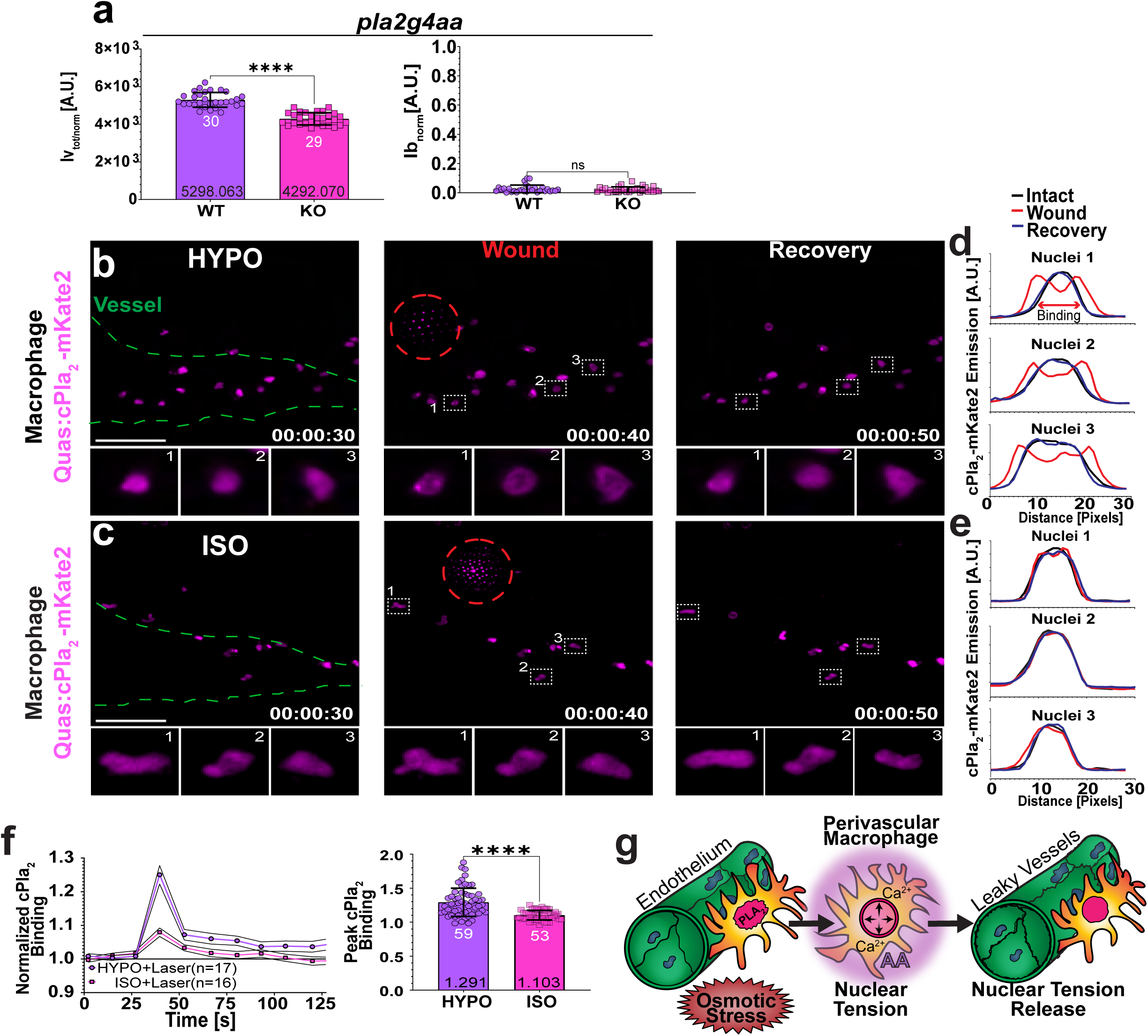
Nuclear shape sensing by cPla_2_ relays rapid osmotic wound cues to vessels. (**a**) Quantification of vessel and wound leakage in homozygous (*pla2g4aa*^mk217/mk217^) and heterozygous (*pla2g4aa*^mk217/wt^ as control) cPla_2_ mutant animals. (**b-c**) Representative MIPs of Tg(*mpeg1.1*: cPla_2_-mKate2) larvae before and after laser injury (dashed, red circle) in HYPO (10 mOsm, upper panel) or ISO (280 mOsm, lower panel) bathing solution. Magenta pseudo-colour, *mpeg1.1*: cPla_2_-mKate2. Green dashed line, vessel outline. Insets, correspond to the cells within the numbered ROIs depicted in the middle panel. Time stamp, hh:mm:ss. Scale bars, 50 μm (**d**) Line profiles of cPla_2_-mKate2 fluorescence in the ROI-labelled cells show translocation after laser injury in HYPO but not in (**e**) ISO bathing solution. (**f**) Left panel, cPla_2_-mKate2-INM binding dynamics. UV laser injury is induced at T= 40 s. INM-binding of cPla_2_-mKate2 is quantified as a ratio of nucleoplasmic to perinuclear fluorescence signal and normalized to its initial (T= 0 s) value. Right panel, comparison of peak translocation. N, number of analysed nuclei from 53 and 59 perivascular macrophages for ISO_(NaCl)_ and HYPO laser wounds, respectively. Note, the panel depicts two different representations of the same dataset. (**g**) Simplified cartoon scheme. Perivascular macrophage nuclei are reversibly stretched by osmotic wound signals. In the presence of Ca^2+^, nuclear membrane tension causes arachidonic acid (AA) release by cPla_2_. AA is metabolized into a vessel permeabilizing lipid mediator. P values, non-paired two-tailed Student’s t-test (**a**) and non-paired two-tailed Mann Whitney test (**f**). White numbers, number of analysed nuclei pooled from 16-17 animals. Black numbers, mean of dataset. Error bars, SD. The figure source data and numerical P values are listed in the Supplementary Excel File 1. ****P ≤ 0.0001. ns, not significant (P > 0.05).

The transgenic larvae were placed in either HYPO or ISO_NaCl_ solution and injured with a microscope-mounted UV-laser during imaging to capture immediate reporter responses by highspeed spinning disk confocal microscopy (**Fig. 4b, c, S5e, g; Supplemental Video 6**). Under hypotonic but not isotonic bathing conditions, laser injury caused cPla_2_-mKate2 translocation to the INM within the entire field of view within ∼ 10 s of injury. Macrophages close (∼ 20 μm) to the laser blast site showed irreversible INM binding and signs of ER fragmentation, as observed for wound margin epithelial cells in our companion paper (Shen et al., 2024). In stark contrast, the cPla_2_-INM interaction was highly transient in distant, perivascular macrophages. Two-photon resonant scanning microscopy confirmed these results at greater time resolution, suggesting that the wound cues required for efficient cPla_2_ translocation, that is, Ca^2+^ and osmotic shock^4, 6, 31^, propagated at a speed of ∼ 50 μm/s through the tissue (**Fig. S6; Supplemental Video 7**). The rapid timing is consistent with ion/small osmolyte diffusion and explains the immediate onset of vessel leakage after osmotic injury (**Fig. 1c, d**).

## Discussion

Zebrafish tail fin wounds cause a rapid efflux of interstitial solutes into the animal’s freshwater environment. With a diffusivity of D_Na+_ ∼ 1.33e^−9^ m^2^/s or D_Cl-_∼ 2.03 e^−9^ m^2^/s, Na^+^ or Clcan diffuse over (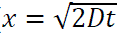) ∼ 50 – 60 μm within just a second. Rapid extracellular ion leakage by the wound triggers an immediate drop of interstitial osmotic pressure along with cell swelling. The cytoplasmic dilution during cell swelling causes colloid osmotic nuclear swelling and INM tension (T_INM_) (Shen et al., 2024), which activates cPla_2_ in synergy with Ca^2+^ ^31^. With some delay, cells then undergo a regulatory volume decrease (RVD) through the expulsion of cytoplasmic solutes and water^38^. cPla_2_ instantly desorbs from the INM, when [Ca^2+^] or T_INM_ drop^6^. Whether the reversibility of cPla_2_-INM interactions reflects RVD or cytoplasmic Ca^2+^ buffering stands to be determined. Compensatory membrane flow from the endoplasmic reticulum (ER) into the contiguous INM may also contribute to T_INM_ dissipation^13^.

Upon injury, perivascular macrophages transiently activate cPla_2_ at distances rarely observed for epithelial cells^4, 6, 13^. Notably, the attribute “perivascular” denotes the predominant macrophage localization (i.e., clustered around the caudal hematopoietic region) in unwounded, larval tail fins— and not (necessarily) a distinct macrophage subtype. Whereas macrophage nuclei must flexibly deform during tissue migration and phagocytosis, epithelial cell nuclei must be more rigid to form stable tissue barriers. Their differences in nuclear mechanosensitivity likely derive from cell-specific, cyto-/ nucleoskeletal requirements and constraints, which determine how much their nuclei can swell. Indeed, F-actin disruption favors nuclear swelling and mechanosensitive cPla_2_ activation after hypotonic shock *in vivo* and *in vitro*^6^. In other words, softer cells (with less cytoskeletal stabilization) may be generally more prone to nuclear membrane mechanotransduction.

When cPla_2_ in perivascular macrophages adsorbs to the INM, it stays there for a few seconds. Our genetic and pharmacologic perturbations suggest that Alox5a and Lta4h metabolize the resulting perivascular AA pulse into a vessel permeabilizing lipid, perhaps LTB_4_^15^. Interestingly, neither macrophages nor vessel leakage are necessary for neutrophil recruitment. However, in line with the known regenerative functions of macrophages^39^, rapid serum exudation may facilitate homeostatic restoration and healing by providing nutrients and growth factors to the stressed tissue.

In summary, our work reveals an essential *in vivo* role for perivascular macrophages as mechanochemical relays for rapid, biophysical wound cues. Whether or how the vascular microenvironment stimulates macrophage mechanosensitivity presents an intriguing question for future research.

## Acknowledgements

We thank Deji Afolalu for aquatic facility management, Mariia Akhmanova and Tasos Siskoglou for their expertise in multiphoton imaging and Denis A. Larochelle, Mariia Akhmanova, Miklós Lengyel and Srivatsan Rajan for valuable comments on manuscript. This research was supported by the NIH/NIGMS grants R35GM140883 to P.N. and Gerstner Sloan Kettering Harold E. Varmus Fellowship (Mr. and Mrs. Rosen Family) to Z.G.

## Author Contributions

PN and ZG conceived the study and designed the experiments. ZG and ZS conducted the experiments. YM, MJ and ZG designed and established the new CRISPR genetic and transgenic lines in the study. ZS and ZG developed the Python and Fiji image analysis scripts and analysed the data. ZG prepared the figures. ZG and PN designed MATLAB scripts for data analysis aided by ChatGPT4.0. ZG and PN wrote the paper.

## Competing interests

Authors declare no competing interest.

## Data and materials availability

Raw data are available from the authors on reasonable request. Analysis scripts and image and data processing pipeline are available on GitHub page for Fiji, Python and MATLAB.

## METHODS

### Zebrafish Husbandry

Adult wild-type and mutant Casper^40^ Zebrafish (Danio rerio) strains and larvae were maintained as described^26^ and subjected to experiments according to the institutional compliance and approval of the animal ethics committee, the Institutional Animal Care and Use Committee (IACUC) and the Research Animal Resource Center (RARC) of the Memorial Sloan Kettering Cancer Center (MSKCC). The adult fish are reared in either 2.8 or 6 L polycarbonate tanks at animal density 8 fish L^−1^, with a photoperiod of 14:10 light:dark cycles and maintained in salinity-conditioned system water surveillance, (pH=7.53) sodium bicarbonate (Proline, SC12A), conductivity 700-850 μS cm^−1^(Instant Ocean Sea Salt (#SS1-160P)) with TGP=99.4% (Total Gas Pressure) and DO=8.18 ppm (Dissolved Oxygen=8.155 mg L^−1^) at 28 **°**C. The zebrafish are fed twice a day with live feed consisting of rotifers or artemia, followed by processed dry spirulina pellets (Zeigler). All anaesthesia was conducted with 0.2 mg ml^−1^ 3-amino benzoic acid ethyl ester (Sigma, MS-222, E10521), (pH=7.0), buffered in 0.5 mg ml^−1^ anhydrous sodium phosphate dibasic (Fisher, BP332-500). The embryos were staged by dpf (days-post fertilization). Sex was indeterminate at 2.5–4 dpf and all experiments in the manuscript were conducted at these embryonic stages. The animal embryos were collected from natural spawning and raised in standard hypotonic E3 containing 0.1%(w/v) methylene blue (Sigma-Aldrich, M9140) for first 24hrs followed by E3 medium (5 mM NaCl (Sigma-Aldrich, S7653), 0.17 mM KCl (Sigma-Aldrich, P9333), 0.33 mM CaCl_2_ (Sigma-Aldrich, C5670), 0.33 mM MgSO_4_ (Sigma-Aldrich, M7506)) in 100 mm petri dishes(Fisher Scientific, FB0875713).

### Transgenesis, plasmid construction and *in vitro* transcription

Fertilized Casper zebrafish embryos were collected and injected at the one-cell stage into the cytoplasm with a laser-pulled borosilicate glass microneedle (H=370, FIL=5, VEL=50, DEL=245, PUL=140; P-2000 Sutter Instrument CO.) and a Nanoject II^TM^ microinjector (Drummond Scientific, Broomball, PA)^41^. Plasmids were assembled using Gateway multisite cloning kit using LR Clonase^TM^ (Invitrogen; C12537-023) and multicloning sites between the gateway att sites flanking cassette of p5E, pME and p3E plasmids that were recombined into Tol2kit^42^ destination vectors pDESTTol2CG2* or pDESTcrybb1 for generating following Q-system animals^43^: Tg(*kdrl*:Qf2)^44^; Tg(*mpeg1.1*:Qf2)^45^; Tg(*lyz*:Qf2)^45^ or Tg(Quas:eGFP)^42^; Tg(5xQUAS:GAP-tagYFP-P2A-NfsB_Vv)^35^ and Tg(Quas:cPla_2_-mKate2-P2A-eGFP-KDEL) or transgenic Casper zebrafish. The latter plasmid was constructed from Cytosolic Phospholipase A2 (cPla_2_, Ensembl: ENSDARG00000024546)^1^ where the open reading frame was amplified by PCR from a zebrafish cDNA clone(Open Biosystems, 9037889) using custom designed primers bearing unique palindromic over-hangs, Bsu15I and EcoRI:(Fwd:5’-AGTCatcgatGCCACAATGTCCAACATTATAG3’; (Rev:5’-CAGTGTGGCTGTGGGAGCTGGAGgaattcGACT-3’). The PCR product was RE digested, gel-purified(Qiagen, 28106) ligated into pDONOR221 containing pME gateway compatible att sites and subsequently fused in-frame to mKate2 (Evrogen) as a C-terminus fluorescent tag with a 15 amino-acid GS-enriched linker^6^. The p3E entry vector was fused in-frame with self-cleaving P2A peptide in N-terminus of eGFP followed by endoplasmic reticulum localization signal KDEL and SV40 poly(A). The final transgenesis constructs were mixed and ∼2.7 nL of 25 ng μL^−1^ of each plasmid co-injected with 25 ng μL^−1^ Tol2kit transposase mRNA, transcribed from NotI linearized pCS2FA-transposase plasmid with mMESSAGE mMACHINE SP6 reverse Transcription kit (Thermo Scientific, AM1340) in wt or Casper embryos. Among the injected embryos, fluorescence-positive siblings in the heart (*cmlc2*:eGFP)^42^, lense (*crybb1*:mKate2)^46^ or chorion (*he1*:eCFP)^35^ were selected and raised in husbandry and backcrossed at sexual maturity to Casper fish. For establishing stable and constitutive tissue-specific fluorescent transgenic zebrafish lines, Qf2 founders were crossed with Quas animals and their progeny were identified through both transgenesis marker and indicated fluorescence in the designated tissue or cell type^43^.

### Reagents, Isotonic reconstitution, and eicosanoid pathway Inhibition

For osmotic surveillance, a battery of iso-osmotic media was constituted by supplementing standard hypotonic E3 media (Π(osmolality)=10 mOsm) with the following osmolytes: 135 mM Na^+^·Cl^−^ (Sigma-Aldrich, S7653); K^+^·Cl^−^ (Sigma-Aldrich, P9333); K^+^·Br^−^ (Sigma-Aldrich, P9881); Na^+^·Br^−^ (Sigma-Aldrich, 220345); Cs^+^·Cl^−^ (Sigma-Aldrich, C3011); Na^+^·C_6_H_11_O_7_^−^ (TCI Chemicals, G0041); C_5_H_14_NO^+^·Cl^−^ (Fisher Scientific, 50213240) and 270mM D-mannitol(Fisher Scientific, BP686), culminating in ISO_(NaCl)_ or ISO* (Π=280 mOsm) E3 media with the indicated reagents^4^. The final osmolality of solutions was approximated with ddH_2_O (18.2MΩ·cm, Elga PURELAB^®^) calibrated handheld optical salinity refractometer (RHS-10ATC Cole-Parmer, EW-81150-31) ∼1.0050:1.025-1.030 *d*^20^ SG Salt^−1^(∼8:35-40 ‰ PPT) for hypotonic (Π=10 mOsm) and all isotonic (Π=280 mOsm) E3 reagents respectively and used for Imaging and analysis. For reconstitution experiments the nucleotides, their derivatives were conducted by ISO_(NaCl)_ to ISO_(NaCl)_ shifting approach, where the latter was constituted with ISO_(NaCl)_ alone as vehicle or supplemented with 5mM following nucleotide biomolecules: adenosine 5’-triphosphate disodium salt hydrate (ATP; Sigma-Aldrich, A26209) and adenosine (Ad; Sigma-Aldrich, A9251) as previously described^5^. The polyunsaturated arachidonic acid (20:4(5,8,11,14-all-cis-eicosatetranoic acid); Sigma-Aldrich, A3611) was administered in parallel to anesthetized and wounded 3dpf Tg(*kdrl*:Qf2:Quas:eGFP) larvae at a final concentration of 5μM in ISO_(NaCl)_ at 5minutes during the imaging experiment. For eicosanoid pathway pharmacological inhibition experiments, Tg (*kdrl*:Qf2:Quas:eGFP) larvae were pre-incubated for 30-45 or 180 min in hypotonic(Π=10 mOsm) E3 supplemented with the following compounds: 50μM Licofelone (Cayman Chemical Company, 100007692), 10μM MK886 (Cayman Chemical Company, 10133) or 130nM Diclofenac (Sigma-Aldrich, 1188800), respectively^29^. The imaging medium also contained the same concentration of the indicated inhibitors dissolved in hypotonic E3 (Π=10 mOsm) and administered 5min after wounding. All water-insoluble biomolecules and inhibitors were dissolved at a maximal concentration of 1% dimethyl sulfoxide, DMSO (Millipore Sigma, 276855) hypotonic (Π=10 mOsm) or ISO_(NaCl)_(Π=280 mOsm) E3 as a vehicle control.

### Larvae preparation, microangiography, and tailfin wounding

Zebrafish larvae (2.5–4 dpf) were screened for transgenes, anesthetized, and oriented with their left lateral side facing up. They were positioned within slots imprinted from a custom-designed mold cast (NIH 3D Print Exchange, ID: 3DPX-021299) in 2% agarose (Fisher Scientific, BP160) dissolved in hypotonic (Π = 10 mOsm) E3 medium. For microangiography, approximately 5 nL of 70 kDa dextran-tetramethylrhodamine (Fisher Scientific, D1818) dissolved in 1× phosphate-buffered saline (PBS; Sigma-Aldrich, 79382) at 1 mg/mL was microinjected into the Common Cardinal Vein (CCV) or the larval atrium, between the gill and pectoral fin. The injection was performed using a borosilicate microneedle (H = 410, FIL = 5, VEL = 40, DEL = 245, PUL = 160; P-2000 Sutter Instrument Co.) and a microinjector (Drummond Scientific, Broomball, PA) under the guidance of a stereomicroscope (Discovery V.8, Zeiss).

Embryos with successful perfusion of fluorescent dextran dyes in the caudal arteries and veins were identified using a coaxial fluorescence dissection stereomicroscope (MVX10, Olympus) equipped with an MVPLAPO 1X objective (NA 0.25, WD 65 mm, FN22) and a mercury lamp (LM200B1-A, Prior Scientific) with dichroic mirror sets (MVX-RFA; 540/35 U-MGFPHQ/XL, 625/55 U-MRFPHQ/XL). For tailfin imaging experiments, anesthetized embryos were flat-mounted on their right side in a 35 mm glass-bottom microscopy imaging dish (MatTek Corporation, P12G-1.5-14F) and immersed in ∼200 μL of 0.8% (w/v) low-melting agarose (Goldbio, A-204-100) dissolved in ISO_(NaCl)_ (Π=280 mOsm) E3medium. For tailfin wounding, multiple embryos were aligned in parallel, and the tips of the larval caudal fins were amputated at the posterior end of the notochord using a tungsten microblade needle (Fine Science Tools, 10318-14). Care was taken to avoid damaging the notochord or caudal vessels during the solidification of the low-melting agarose^4, 24^. After immobilization, the embedded live larvae were enveloped in 200 μL of ISO_(NaCl)_ (Π=280 mOsm) containing tricaine to prevent desiccation. The immobilized larvae are allowed to rest at 28 °C for approximately 15 minutes to restore cardiac function in the trunk. Following this, the tailfin region of the embedded larvae was carefully freed from the excess low-melting agarose using a tapered micro spatula (Fine Science Tools, 10089-11) and transferred to a pre-heated stage-top incubator (INU-TIZ-D35).

### Spinning disk confocal microscopy

Tailfin amputation Imaging experiments were completed at 28 **°**C with the heated imaging chamber (INUG2 KIW, TOKAI-HIT) and sub-stage heater (AirTherm ATX In vivo Scientific, WPI inc.), in an inverted Nikon Eclipse Ti microscope equipped with a CFI Apo LWD Lambda S-series 20X Objective lens (N.A.=0.9 ꚙ/0.11-0.23 WD=0.95μm WI Objective, Nikon). The samples were excited with 488 and 561 nm diode laser (LUN-F 100-240V∼50/60Hz, Nikon Instruments inc.), where channel acquisition intensities/exposure times used in the manuscript were as follows: 40%/100ms (488 nm, 30 mW or 0.03 VA) or 25%/100ms (561 nm, 11.5 mW or 0.015 VA) laser power settings for illuminating either endothelium or innate immune cells(macrophages or neutrophils) and Dextran, respectively. The fluorescence emission spectra were collected in 4D(XYZT) by automotive PCI hardware triggering (PXI-1033, National Instruments) with a Yokogawa CSU-W1 Spinning Disk unit (Nipkow disk pinhole=50μm, 4000rpm), incorporating a Photometrics Prime BSI Scientific CMOS (sCMOS) camera with analog Z-plane acquisition (MCL Nano-Drive NIDAQ Piezo Z, Nikon) and a motorized XY-stage (TI-S-ER, Nikon). The emission was collected for either green transgenic (Ex/Em:488/525; Gain: Correlated Multi-Sampling) or red Dextran (Ex/Em:561/605; Gain: High Dynamic Range) with a high-sensitive sCMOS camera using band-pass dichroic mirrors(Chroma Technology Corp., 89100bs) for filtering two separate fluorescence emission spectra (525/36 and 605/52, Chroma Technology Corp., 89000 Sedat Quad) placed in front of the detector for isolated detection of green and red fluorescence, respectively. The imaging plane (1331 µm x 1331 µm x 100 µm) was collected at 3 µm z step size to result with 35 slices and repeated with 30-second intervals per position for up to 60 minutes. Collectively, the live embryos were exposed to 5-6.81 W cm^−2^ and 2-3.1 W cm^−2^ laser power density for 488 and 561nm, respectively. For all imaging experiments, the imaging dish was covered with a solution of interest at 270-300s (T=10-11^th^ frame) with a bolus of 10x agarose volume(∼2mL). The z-stack images of zebrafish embryos were captured using multiple scanning modes at 100µm tissue depth in z-axis at a resolution of 2048 × 2048 pixels (16 bit) in the x and y plane, corresponding to 0.645 µm/pixel calibration with a voxel size of (0.65 × 0.65 × 3 µm) in x y and z, respectively. The high-resolution XYZ-T image files were pre-processed by (2×2) binning and acquired as (1024 × 1024 per z-plane) with corresponding 1.33 µm/pixel calibration through triggered excitation and emission collection for either red or green fluorescence in the NIS imaging software (NIS Elements, 5.12.0). Under these microscope settings, endothelial responses were imaged with a high spatiotemporal resolution where each 3D stack (frame) took approximately 14 seconds (4 Z-stacks per minute) allowing multi-position acquisition of two transgenic or four dextran-perfused larvae in parallel in a single imaging experiment.

### Spinning disk confocal microscopy and laser Wounding

For laser wounding experiments, single intact and anesthetized 3 dpf Tg (*mpeg1.1*:Qf2;Quas:cPla_2_-mKate2-P2A-eGFP-KDEL) were embedded in 60mm plastic petri dish (Corning, 351007) and immobilized with ∼200μL 1% LM agarose ISO_(NaCl)_ (Π=280 mOsm) E3. Subsequently, the LM-agarose was enveloped with ∼2-3 mL of hypotonic (Π=10 mOsm) or ISO_(NaCl)_(Π=280 mOsm) E3 media for creating a submerging environment for the 25X Objective lens (N.A.=1.1 ꚙ/0-0.17 WD=2μm Water Dipping, Nikon) in Nikon Eclipse FN1 upright microscope. The samples were excited with 488 and 561nm diode laser lines (Andor Revolution XD) and fluorescence channel excitation intensity/exposure was adjusted to 35%/80ms(488nm) and 30%/80ms(561nm) for illuminating macrophage endoplasmic reticulum (KDEL-eGFP) and cPla_2_-mKate2 in the caudal region of the larvae. The emission spectrum were excited in 4D(XYZT) by SmartShutter controller (Sutter Instrument, LB10-B/IQ Lambda) triggered excitation carried by Andor Laser combiner (LC-501A, Andor Technology) housing Andor iXon3 897 thermoelectrically cooled EMCCD camera with analog Z-plane acquisition (STG-STEPPER-Piezo Focus, Ludl Electronic Products). The emission was collected with a Yokogawa CSU-X1 spinning disk unit (Nipkow disk pinhole=50μm, 10,000rpm) for green (Ex/Em:488/525) or red (Ex/Em: 561/625) in electron-multiplying mode(Gain: 10MHz at 14-bit; Multiplier 300, conversion 1.0X) and band-pass filters for green (525/40, Semrock., FF02-525/40-25) or red (617/73, Semrock., FF02-617/73-25) fluorescence emission spectra. The imaging plane (287 µm X 287 µm X 50-80 µm) was collected at 1.5µm z step size and repeated in no-delay intervals per position for up to 30 minutes, resulting with temporal step resolution of ∼7-12 seconds per stack. The z-stacks were acquired at a resolution of 512 × 512 pixels (14-bit), corresponding to 0.561 µm/pixel calibration with a voxel size (0.56 × 0.56 × 1.5 µm) in x y and z, in the NIS imaging software (NIS Elements, 3.22.14). Collectively, live unwounded embryos were exposed to 5 W cm^−2^ and 4 W cm^−2^ laser power density for 488 and 561nm, respectively. The wounds were induced at ∼1 minute(T=5-7), with several successive laser pulses targeted at peripheral sentinel macrophages in a single frame with 60ms delay using micro-scope-mounted 435nm ultraviolet Micropoint Laser (Andor), resulting in tailfin fin fold ablation around the boundary of hematopoietic region in the trunk of the larvae.

#### Multiphoton imaging and FRAP-wounding

3dpf intact larvae Tg(*mpeg1.1*:Qf2;Quas: cPla_2_-mKate2-P2A-eGFP-KDEL) embryos were mounted in 35mm glass-bottom microscopy imaging dish (MatTek Corporation, P12G-1.5-14F) and transferred to Ti2-E inverted Nikon microscope, housing AX-R multiphoton modality(AX-MP NDD) and stage top incubator set to 28 **°**C (TOKAI-HIT, STX). The multiphoton images were excited through simultaneous absorption of 920nm tuneable and 1045nm fixed two photon lasers (mks, spectra-physics) for eGFP and mKate2, respectively. The excitation was adjusted to 920nm (4.3%, Gain=45, Line Averaging=4X) and 1045nm (5.3%,Gain=45, Line Averaging=4X) for eGFP and mKate2, respectively. The fluorescence emission was collected using 40X (Plan Apo lambda S 40XC, NA=1.25 ꚙ/0.13-0.21WD=300μm, Nikon) silicone objective with objective heating mantle (28 **°**C) and detected with PMT GaAsP. The bidirectional images were acquired in resonant scanning mode. The imaging plane (147 µm X 90 µm X 15 µm) is collected at Nyquist optical resolution of 0.149 µm at 1.5 µm z-step size using NIDAQ piezo and repeated in no-delay intervals for up to 2 minutes, resulting with a temporal step resolution of 2.5 seconds per stack. The z-stacks were acquired at a pixel resolution of 1024×626 (14-bit), corresponding to 0.144 µm/pixel calibration with a voxel size (0.14 × 0.14 × 1.5 µm) in x y and z in the NIS imaging software (NIS elements, 6.02.01). The wounds were induced at ∼1 minute with a single high intensity 920nm laser pulse (22%, Line Averaging=1X, Dwell time=1s, area= 20 × 20 µm) with photostimulation MP STIM FRAP modality in Galvano scanner mode, resulting in epithelial tailfin fold laser ablation around the boundary of hematopoietic region in the larvae.

#### Denoise and 3D Deconvolution

The 4D image stacks of tg(*mpeg1.1*:Qf2;Quas: cPla_2_-mKate2-P2A-eGFP-KDEL) were corrected by 3D deconvolution, where sample PSF was derived and computed automatically with depth-calibration and without image intensity subtraction or preprocessing in Nikon NIS elements (5.21.03, Build 1489). The deconvolution process was carried by default method using landweber algorithm for spinning-disc modality with 50 µm pinhole size with immersion refractive index 1.33 (water) for both 488 and 561nm fluorescent light emission^47^. The deconvolved or MP image stacks are denoised using denoise.ai tool for improving signal-to-noise in Nikon NIS elements (5.21.03).

### Generation of zebrafish CRISPR mutants

To generate zebrafish mutants a CRISPR/Cas9 system with a single sgRNA (Alox12-sgRNA1) targeting zebrafish *alox12* (ENSDARG00000069463) exon7 is used. The ribonucleoprotein complex consisting of *alox12*-sgRNA1 and Cas9 recombinant protein is injected into the cytoplasm of one-cell stage fertilized zebrafish embryos. The injected F0 larvae were brought to adulthood and crossed with wild-type adults to produce F1 progeny. The *alox12*-sgRNA injected F1 zebrafish were grown to sexual maturity and their genomic DNA was isolated from their tail fins for genotyping. Tail fins were partially amputated, suspended in 250 μL of 0.05M NaOH, incubated at 95**°**C for 10 minutes, cooled on ice for 10 minutes, then neutralized with 25 μL of 1 M Tris-HCl (pH 8), and vortexed. The DNA extracts were used as genomic templates for Polymerase Chain Reaction (PCR). 486bp PCR products were digested with FastDigest BseLI (Thermo Fisher Scientific, FD1204) overnight at 37**°**C, producing two DNA fragments (310 and 176bp). The 508bp represents a mutant allele where the BseLI site has been disrupted by Cas9-induced mutation. The 508bp band was isolated from the agarose gel and sequenced via sanger, confirming a 22bp insertion (**Fig. S2 I, j**). Subsequently, the F1 heterozygous adult zebrafish with 22bp frameshift mutation were bred to homozygosity.

Two independent sgRNAs (*lta_4_h*-sgRNA-1 and *lta_4_h*-sgRNA-3) targeting zebrafish *lta_4_h* (ENSDART00000028171) were used to establish a CRISPR-zebrafish line, as previously described^24^. The Cas9-gRNA ribonucleoprotein complex (a combination of *lta_4_h*-sgRNA1 and *lta_4_h*-sgRNA3) was injected into the cytoplasm of one-cell-stage zebrafish embryos. After the injected F0 larvae matured (2-3 months post-fertilization), individual F0 adults were crossed with wild-type adults to produce F1 progeny. These F1 larvae were then grown to sexual maturity, and genomic DNA was isolated from their tail fins for genotyping. For *lta_4_h*-sgRNA1, PCR products were digested with FastDigest BseLI (Thermo fisher Scientific; FD1204), producing three DNA fragments (271, 140, 129 bp). For the *lta_4_h*-sgRNA3, PCR products were digested with FastDigest SalI (Thermo fisher Scientific; FD0644), producing three DNA fragments (271, 184, 85 bp). The 271 bp product represents a mutant allele where the BseLI or SalI site has been disrupted by Cas9-induced mutation. This 271 bp band was isolated from the agarose gel and sequenced via Sanger sequencing, confirming a 5 bp deletion (2, 2, 1 bp deletion) and a 7 bp insertion (**Fig. S2k, l**). F1 heterozygous adult zebrafish with this frameshift mutation of interest were bred to achieve homozygosity.

A single guide was designed targeting zebrafish *pla2g4aa* (ENSDARG00000024546) for genetic disruption at exon 7. The Cas9-gRNA ribonucleoprotein complex consisting of *pla2g4aa*-sgRNA1 was injected into the cytoplasm of one-cell-stage fertilized embryos. The injected larvae were then brought to adulthood and the individual F0 animals were used to produce F1 progeny by backcrossing with wild-type Casper fish. The F1 larvae were grown to sexual maturity, genomic DNA was subsequently isolated from their tail fins and sequenced. The *pla2g4aa*^wt/mk217^ DNA was PCR amplified, digested with FastDigest BseMI (Thermo fisher Scientific; FD1264) and then analysed by agarose gel electrophoresis for genotyping. The 447bp PCR product from the wt *pla2g4aa* allele is cleaved into two smaller products upon overnight incubation with BseMI (281 and 166bp) while the 437bp PCR product of the mk217 allele is not cleaved by restriction digest into the smaller products. F1 heterozygous adult fish with the mk217 10bp deletion (**Fig. S5a, b**) were bred to achieve homozygosity.

Alox12 (Chr17, *alox12* Ensembl: ENSG00000108839) mk215 allele Exon7-ins-22bp

gRNA2: GGATCACTGGGCAGAAATAC TGG

Fwd: 5’-CAAATGCATTGATGCAAAAAGT-’3

Rev: 5’-TGAGAAATAGCATTCATTTGCG-’3

Lta_4_h (Chr 12, *lta_4_h* Ensembl: ENSG00000111144) mk216 allele Exon1-ins-7bp-Δ5bp

gRNA1 GAAAGTCGCCCTGACTGTGG AGG

gRNA3 CATGCCTGTCAAAGTCGACA TGG

Fwd: 5’-TCAACCATGACTCCAGTTTCAG-’3

Rev: 5’-CAGTGCATTGGATCGTACTCAT-’3

cPla_2_ (Chr1, *pla2g4aa* Ensembl: ENSDARG00000024546) mk217 allele Exon7Δ10bp

gRNA1 GGAGGTTTTCGTGCAATGGT GGG

Fwd: 5’-CCATCCTCACCAAGAGAGGTAA-’3

Rev: 5’-ACTGCTTGAATTGACTGCAAAA-3’

### Larval genotyping of established zebrafish CRISPR mutants

Zebrafish Larvae were recovered from the 35mm imaging dish and subjected to 20μL 0.05M NaOH alkaline DNA extraction at 95**°**C and neutralization with 2 μL 1M Tris-HCl (pH 7.4)^24^. The gRNA targeted exon was PCR amplified and subjected to restriction enzyme digestion followed by gel electrophoresis analysis. For imaging experiments where the genotype was unknown (in-cross of heterozygotes for pla2g4aa), blinding was inherent in the experimental design and the genotypes were matched to the assigned fish *post hoc*. The previously published zebrafish CRISPR larvae were genotyped using the following primers:

Alox5 (Chr13, *alox5a* Ensembl: ENSDARG0000057273) mk211allele Exon7Δ10bp^29^

gRNA1: TGGGTGCCGCCAAGTACTGA TGG

Fwd: 5’-GCTGTAATCCAGTGGTCATCAA-’3

Rev: 5’-TGATCTCACTGGAGACTGGAGA-’3

#### Chemogenetic depletion and Sudan Black staining

To perform macrophage-specific cell depletion, Tg(*mpeg1.1*:Qf2;5xQuas:GAP-tagYFP-P2A-NfsB_Vv) animals tagged with nitroreductase under the mpeg promoter were used. Tg(*lyz*:Qf2;5xQUAS:GAP-tagYFP-P2A-NfsB_Vv) was used as a genetic vehicle control. The 100 mM MTZ (Sigma-Aldrich, 1442009) stock solution was prepared in DMSO (Millipore Sigma, 276855) and protected from light. Embryos were dechorionated prior to metronidazole (MTZ) ablation using 1 mg/mL pronase (Roche, 165921). Starting at 2 dpf, the larvae were treated with 150 μM MTZ and 0.15% DMSO in hypotonic (Π = 10 mOsm) E3 media or mock-treated with 0.15% DMSO in hypotonic E3 as a vehicle control for at least 24 hours. At 3 dpf, MTZ was temporarily removed and replaced with hypotonic (Π = 10 mOsm) E3 media until the microangiography procedure and subsequent immobilization in ISO_(NaCl)_(Π = 280 mOsm) low-melting agarose for confocal imaging. During confocal imaging, larvae were shifted back to hypotonic (Π = 10 mOsm) E3 media containing either 150 μM MTZ or 0.15% DMSO, depending on their respective treatment conditions. For pharmacological inhibition of eicosanoid pathway, macrophage depleted or DMSO treated larvae were both inhibited with 30-45 min 50μM Licofelone dissolved in hypotonic (Π=10 mOsm) E3 and subjected to dextran microangiography and imaging. For Sudan Black staining, mpeg-depleted or DMSO vehicle larvae were wounded and fixed 90min post wounding using 4% paraformaldehyde (Fisher Scientific, BP531) in 1x PBS overnight at 4 **°**C and then stained with Sudan-Black for 30 min. The larvae were rinsed three times in 70% ethanol (Decon Labs, 2405) and rehydrated with PBS-Tween-20 for 5 min. Prior to imaging, the larvae were depigmented(1% KOH, 1% H_2_O_2_) and washed for total of three times in PBST solution 5 min each and transferred for transmitted light imaging^24^.

#### Image processing and measurement of interstitial serum leakage and Dilation

Nikon ND2 Spinning disk confocal time-lapse stacks of wounded larvae were imported into Fiji (v1.54J Just ImageJ Package) using bio-formats plugin and z-projected (Maximum-Intensity Projection) with brightness and contrast-enhanced. To minimize the movement of the imaged plane, the 3D(XYTZ=MAX) was registered using 2D Linear Stack Alignment SIFT multichannel (PTBIOP plugin) tool and the dextran channel was set as reference on the imported composite image stacks (Scale Invariant Interest Point Detector) and rigid transformed (maximal alignment error:50px, inlier ratio:0.05) for subsequent morphology or intensity analysis of endothelium or and dextran respectively. For genetic experiments, the registration was carried out using SIFT Linear Stack Alignment with default settings. For quantification of serum leakage, dextran intensity was traced per embryo in a stratified manner in the extra-epidermal 20-50 μm wounded region (I_bleed_(t)) and below the caudal arteries & veins (I_vessel_(t)). Both Iv and Ib datasets were subtracted with background (empty area in field of view) intensity per timepoint for correcting the noise of sCMOS camera. The quantifications were carried out manually to monitor the accuracy of defined regions in the tailfin using set rectangle tool with fixed dimensions for wound (50 × 50 pixels), vessel (300 × 100 pixels), respectively and quantified using plot z-axis profile function. For endothelial morphological parameter measurements and dilation, vertical line was drawn using line tool with fixed dimensions overlaying the split and registered endothelial fluorescence signal channel (length=50 pixels, with=30) and multi-kymograph function with default settings. The resulting kymograph was binarized (Threshold method=Otsu, Dark Background) to create an 8-bit image and area quantified using makeline tool (width=30 pixels, length=121) and the diameter measured using the plot profile function. All imaging data was normalized to the first 10 timepoints per imaged embryo using custom Python (Conda, v3.8.10 64-bit) and MATLAB (R2024a, v24.1.0.2537033, 64-bit) code and combined (Github: https://github.com/joeshen123/Nuclear-membrane-analysis-in-zebrafish-macrophage; https://github.com/zazadovv/zazadovv). The samples that have shifted out of the imaging plane during the course of experiment were excluded from the analysis. The images’ dimensions are set to be consistent between all samples, and these measurements are designated and independently represented.

#### Image-based measurement of cPla_2_ nuclear membrane binding in macrophages

The 4D spinning disk confocal stacks were imported into napari (v0.4.10) and the cPla_2_-mKate2 channel was segmented in 3D by isolating the middle frame based on the contour area and seeded for watershed segmentation. The background was subtracted in z-axis using rolling ball function and the non-track objects were cleared and tracks of segmented image stack were generated by defining mid-section area, volume and protein binding by defining properties of middle frame area and cPla_2_-mKate2 channel intensity. The obtained 3D tracks were directed to normalization and edge-preserving smooth function. The obtained tracks were quantified for nucleus volume geometry and protein binding over time and the discontinuous tracks were excluded from the analysis. all frames among different remaining tracks were unionized and normalized to cPla_2_-mKate2 intensity in the 1^st^ frame of the image stack and quantified as membrane binding.

#### cPla_2_-mKate2 emission and 2-D quantification

The Tg(*mpeg1.1*-cPla_2_-mKate2) emission was quantified for signal distribution in 2D using Fiji (v1.54J Just ImageJ Package). The quantifications were conducted on isolated macrophage Images that were MIP processed and duplicated from whole field view stacks (XY-T) and registered using SIFT Linear Stack Alignment with default settings. The obtained images were quantified using “line” tool and “Plot Profile” function by drawing line-plots on the periphery of macrophage nuclei and quantifying its distribution over set time-points. The obtained data were then plotted for analysis of transient cPla_2_-mKate2 localization.

#### Quantification of leukocyte recruitment

The still images of the Sudan-black stained larvae were processed in Fiji with z-project function (Maximum-Intensity Projection), duplicated and blurred (Gaussian, sigma=50 scaled). The background subtracted using image calculator function (“Subtract create 32-bit”, “”), resulting 32-bit floating image was median-filtered (radius=5) and quantified for number of leukocytes in the tailfin using find maxima tool (prominence=1950 strict light) and counted. The counts were verified for accuracy by manual counting and analysed.

#### Statistics and Reproducibility

No statistical methods were used to predetermine sample size and the sample size is similar to those reported in previous publications^4, 29^, in brief we always used sample sizes of 8 different animals or more in the study, for laser wounding 3 different animals or more. Unless otherwise indicated, the data analysis was performed blind to the condition of the experiments and each experiment was repeated at least twice with similar results. The experiments were not randomized, and investigators were not blinded to allocation except for genetic experiments with *pla2g4aa* where all the measurements were taken prior to the authors knowing the genotype status of the embryos. The data was sorted into groups post genotyping. Blinding was not performed for the sake of increasing experimental throughput. The calculations in Fiji and napari were completed before calculations in MATLAB and significance quantification in Prism. The statistical analysis was completed using GraphPad Prism 10 (version 10.2.3). Each dataset was tested for normal distribution (Shapiro-wilk or D’agostino-Pearson) test. Only if the data were normally distributed, a parametric method (unpaired two-tailed students t-test) was applied to the dataset. A non-parametric test (two-sided Mann-Whitney) was applied for non-normally distributed data sets. In case of multiple comparisons, one-way ANOVA or Welch one-way with Dunnett’s multiple comparison post-hoc test applied. For analysis with different variance in error distribution, Welch’s correction was used Data from two groups were compared using two-tailed unpaired t-test. *P* values <0.05 were considered significant. Data were represented as box plots where one dot in a graph represents n=1 biologically independent larvae experiment (single embryo). Dot plot Annotated numbers represent the total *n* per group at the top and mean value in the bottom of the graph. In box plots top and bottom edges indicate SD range and midline central marking is representative of mean values. The Data distributions errors are shaded and represented as 95% confidence intervals, unless otherwise indicated. *P<0.05, **P<0.01, ***P<0.001, ****P<0.0001 and ns, not significant. For every treatment, treated and control embryos were derived from the same spawn of eggs. Animals were never reused for different experiments. The larvae were selected on the following criteria: normal morphology, a beating heart and circulating red blood cells. All images shown in the figures are representatives of the respective phenotypes and expression patterns.

**Figure S1. Extended data supporting figure 1.**
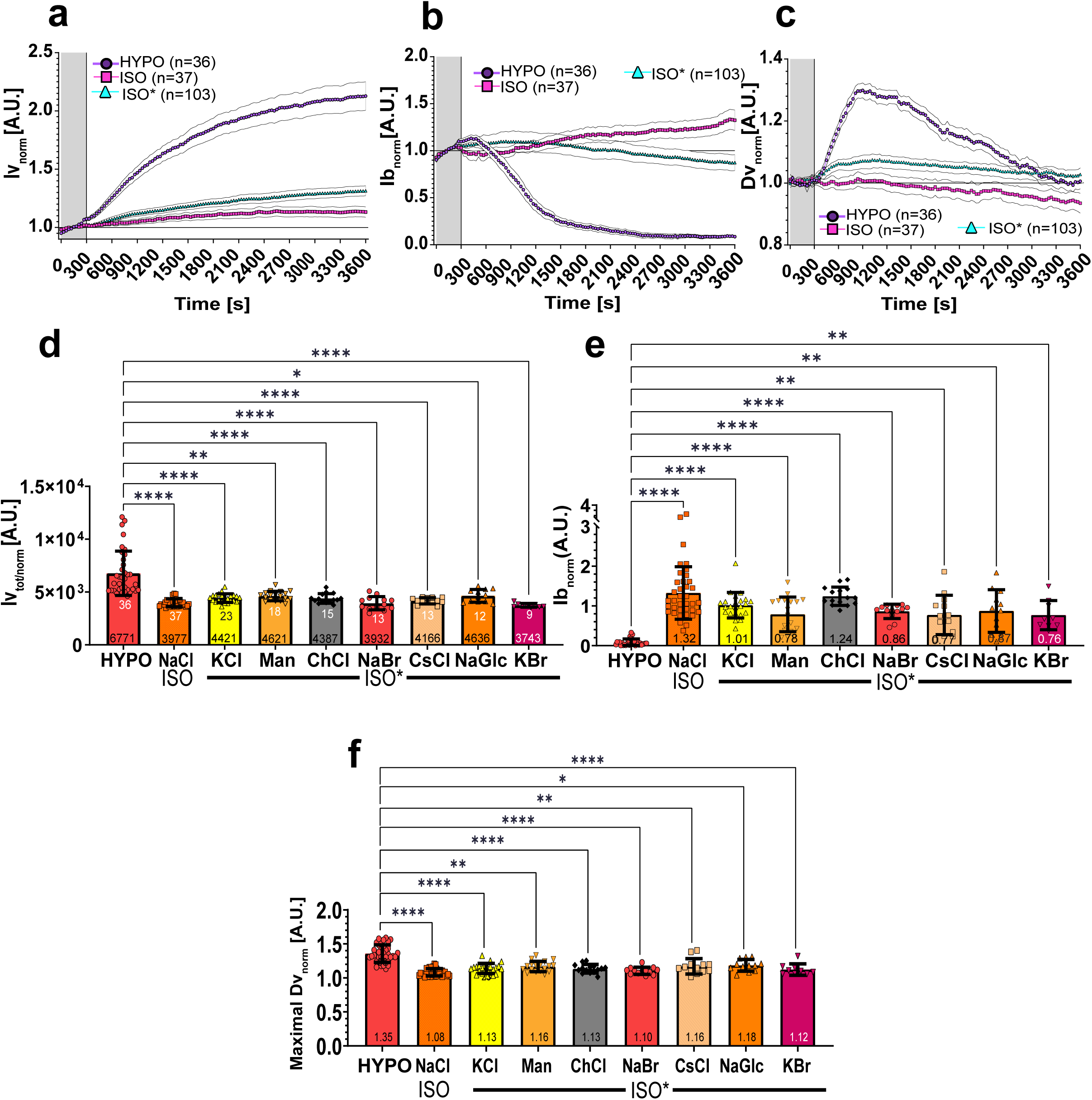
Dynamics of normalized. (**a**) vessel leakage (Iv_tot/norm_(T)), (**b**) wound leakage (Ib_norm_ (T)), and (**c**) vessel dilation (Av_norm_(T)) at the indicated conditions. HYPO, regular E3. ISO, 280 mOsm NaCl in E3. ISO*, pooled isotonic treatments: KCl, potassium chloride. Man, mannitol. ChCl, choline chloride. CsCl, caesium chloride. NaGlc, sodium gluconate. KBr, potassium bromide. NaBr, sodium bromide. The data was normalized to the mean of the first ten frames (grey shaded bar, T= 0-270 s). Error margins, 95% confidence interval of the indicated numbers of animals. Plots of normalized, (**d**) integrated vessel leakage (T=0-3600 s), (**e**) steady state wound leakage (T=3600 s), and (**f**) maximal vessel dilation at the indicated osmolyte conditions. White numbers, animals. Bottom of bar graph numbers, mean of dataset. P values were determined using ANOVA with Dunn’s post-hoc. All figure source data and numerical P values are listed in the Supplementary Excel File 1. *P ≤ 0.05, **P ≤ 0.01, ***P ≤ 0.001, ****P ≤ 0.0001. ns, not significant (P > 0.05).

**Figure S2. Extended data supporting Fig. 2.**
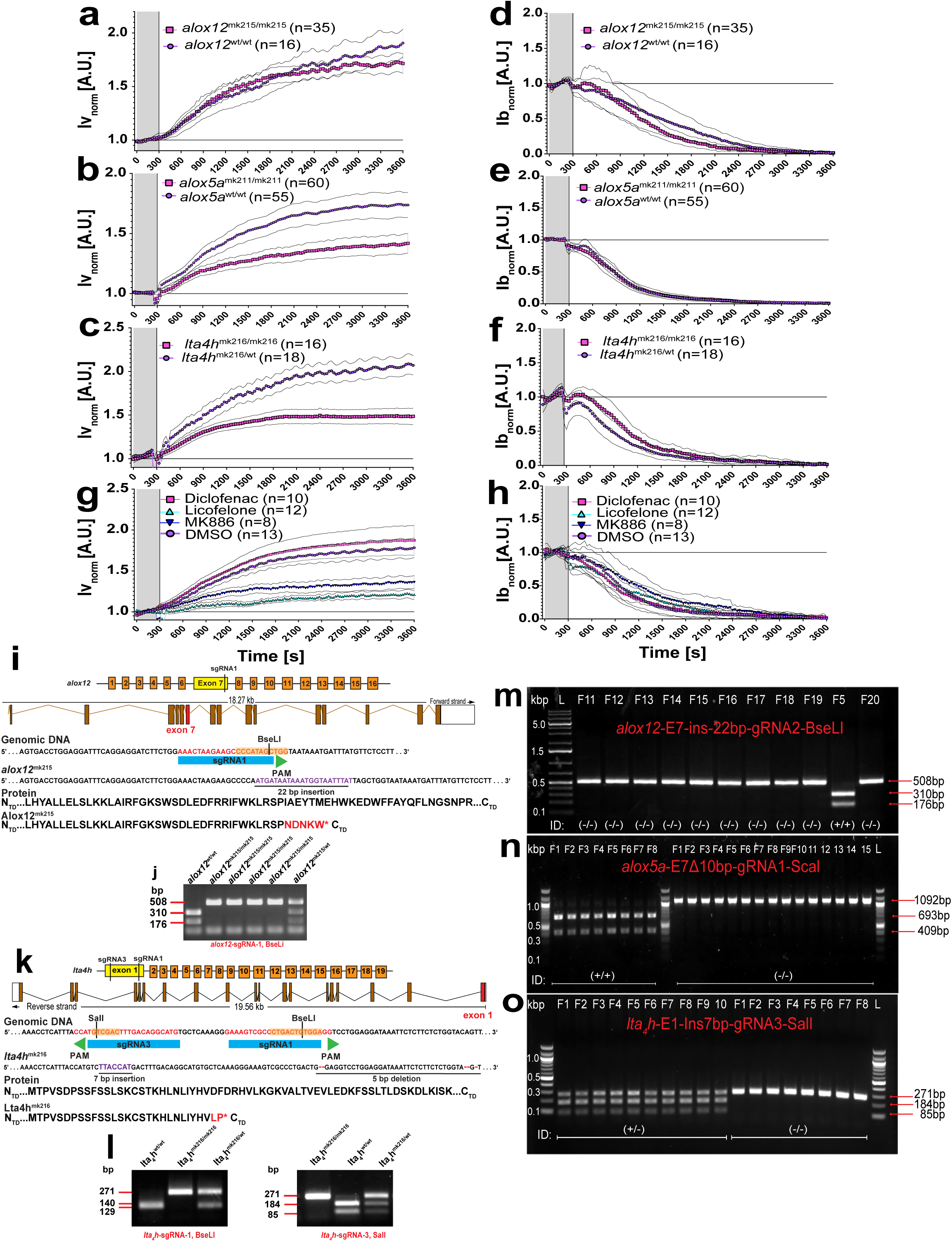
Normalized. (**a, b, c, g**) vessel-(Iv_norm_(T)) and (**d, e, f, h**) wound-(Ib_norm_ (T)) leakage dynamics after ISO_(NaCl)_ → HYPO shifting at the indicated genetic and pharmacologic conditions. (**i**) Scheme illustrating the design guide RNAs targeting exon 7 of alox12. The architecture of the exons (brown and red) and introns (black lines) in the alox12 genomic region. The successful gene disruption is validated by BseLI (CCCATAG^CTGG) exon 7 restriction sites (yellow box) on genomic DNA, which is altered in *alox12*^mk215/mk215^ mutants. (**j**) 486 bp polymerase chain reaction (PCR) product of the wildtype allele is cleaved into two fragments (310, 176bp) by BseLI. The mk215 allele carries a genomic 22bp insertion, preventing the targeted BseLI restriction enzyme digest in *alox12*^mk215/wt^ (bp= 508, 310, 176) and *alox12*^mk215/mk215^ (bp=508). (**k**) Scheme illustrating the design of two single guide RNAs targeting exon1 of *lta4h*. The architecture of exons (brown and red) and introns (black lines) in the *lta_4_h* genomic region is shown. Successful gene disruption is validated by BseLI (CTGACTG^TGGA) and Sall (G^TCGAC) exon1 restriction sites (yellow box) on genomic DNA, which is altered in *lta_4_h*^mk216/mk216^ mutants. (**l**)The 269 bp polymerase chain reaction (PCR) product of the wildtype allele is cleaved into two BseLI (140 and 129 bp) or Sall (184 and 85 bp) fragments. The mk216 allele carries a genomic 7bp insertion and 5bp deletion, preventing the targeted SalI or BseLI restriction enzyme digest in *lta_4_h*^mk216/wt^ (bp=271, 140 129; 271, 184 85) and *lta_4_h*^mk216/mk216^(271bp). The F0 heterozygotes were identified by the presence of three gel bands corresponding to PCR product enzyme digest of adult tailfins. (**m-o**) Mutations were validated for dextran injected larva after imaging. Representative ethidium bromide DNA with set PCR product restriction enzyme digest with BseLI (bp=310, 176) for *alox12*^mk215/mk215^ (bp=486) and *alox12*^wt/wt^ (bp=310, 176) (**m**), ScaI (bp=693, 409) *alox5a*^mk211/mk211^ (bp=1102) and *alox5a*^wt/wt^ (bp=693, 409) (**n**) and SalI (bp=184, 85bp) *lta_4_h*^mk216/mk216^ (bp=271) and *lta_4_h*^mk216/wt^ (bp= 271, 184, 85) (**o**), where each well corresponds to a single imaged and analysed larva.

**Figure S3. Extended data supporting Fig. 2.**
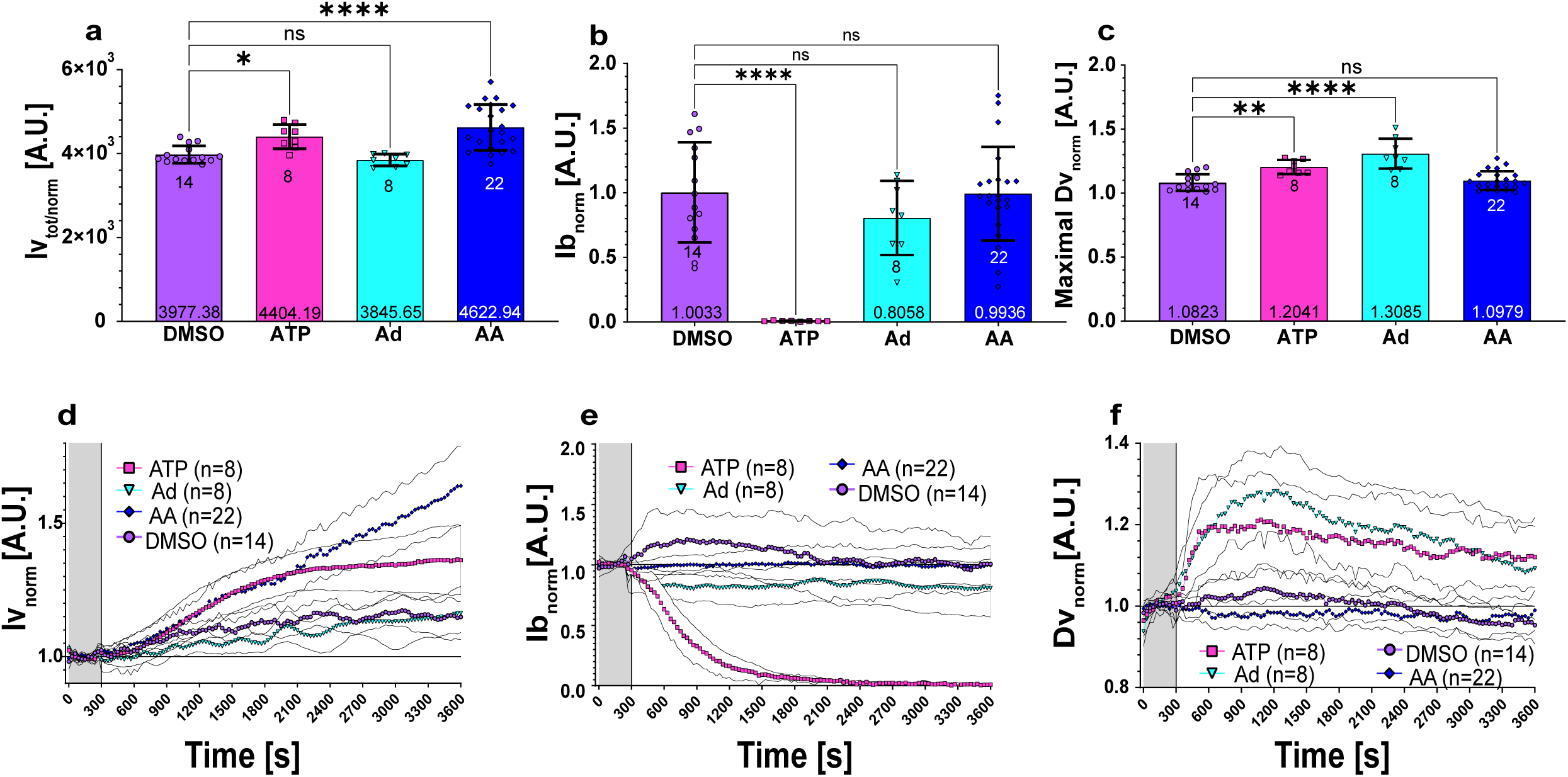
Plots of normalized, (**a**) integrated vessel leakage (T=0-3600 s), (**b**) steady state wound leakage (T=3600 s), and (**c**) maximal vessel dilation upon treatment of wounded zebrafish larvae bathed in ISO_(NaCl)_ with the indicated agonists. DMSO, vehicle. ATP, adenosine triphosphate (5 mM). Ad, adenosine (5 mM). AA, arachidonic acid (5 μM). numbers below the error bars, animals. numbers at the bottom of bar graph, mean of dataset. Error bars, SD. Normalized (**d**) vessel leakage (Iv_norm_(T)), (**d, e, f, h**) wound leakage (Ib_norm_ (T)), and vessel dilation dynamics after ISO_(NaCl)_ → HYPO shifting with the indicated agonists. Error margin, 95% confidence interval. Note, (a-c) and (d-f) refer to the same data set. P values, one-way ANOVA with Dunnett’s post-hoc significance tests. All figure source data and numerical P values are listed in the Supplementary Excel File 1. *P ≤ 0.05, **P ≤ 0.01, ***P ≤ 0.001, ****P ≤ 0.0001. ns, not significant (P > 0.05).

**Figure S4. Extended data supporting Fig. 3.**
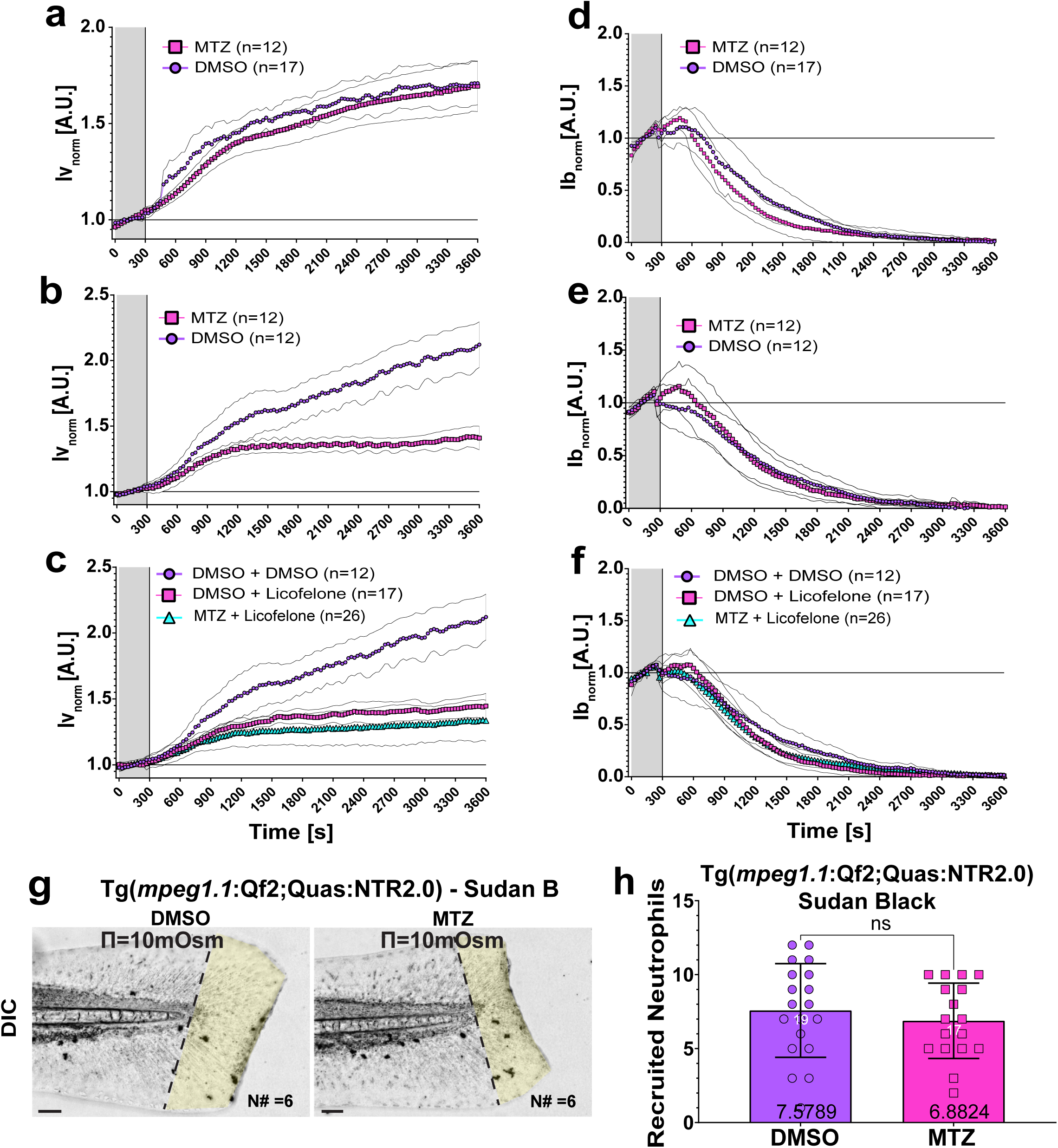
Normalized (**a-c**) vessel-(Iv_norm_(T)) and (**d-f**) wound-(Ib_norm_ (T)) leakage dynamics after ISO_(NaCl)_ → HYPO shifting at the indicated pharmacologic conditions. Error margin, 95% confidence interval. (**g**) Representative transmitted light image of fixed, control (DMSO) and macrophage depleted (MTZ) tail fins of *mpeg1.1*:Qf2;Quas:NTR2.0 larvae at 90 min post injury. Neutrophils are stained with Sudan Black. N#, number of recruited neutrophils in the tailfin region from notochord to the wound edge (highlighted in yellow). Scale bars, 50 μm. (**h**) Quantification of Sudan Black staining. White numbers, animals. Black numbers, mean of dataset. P values, nonpaired two-tailed Student’s t-test. The figure source data and numerical P values are listed in the Supplementary Excel File 1. Ns, not significant (P > 0.05). Error bars, SD.

**Figure S5. Extended data supporting Fig. 4.**
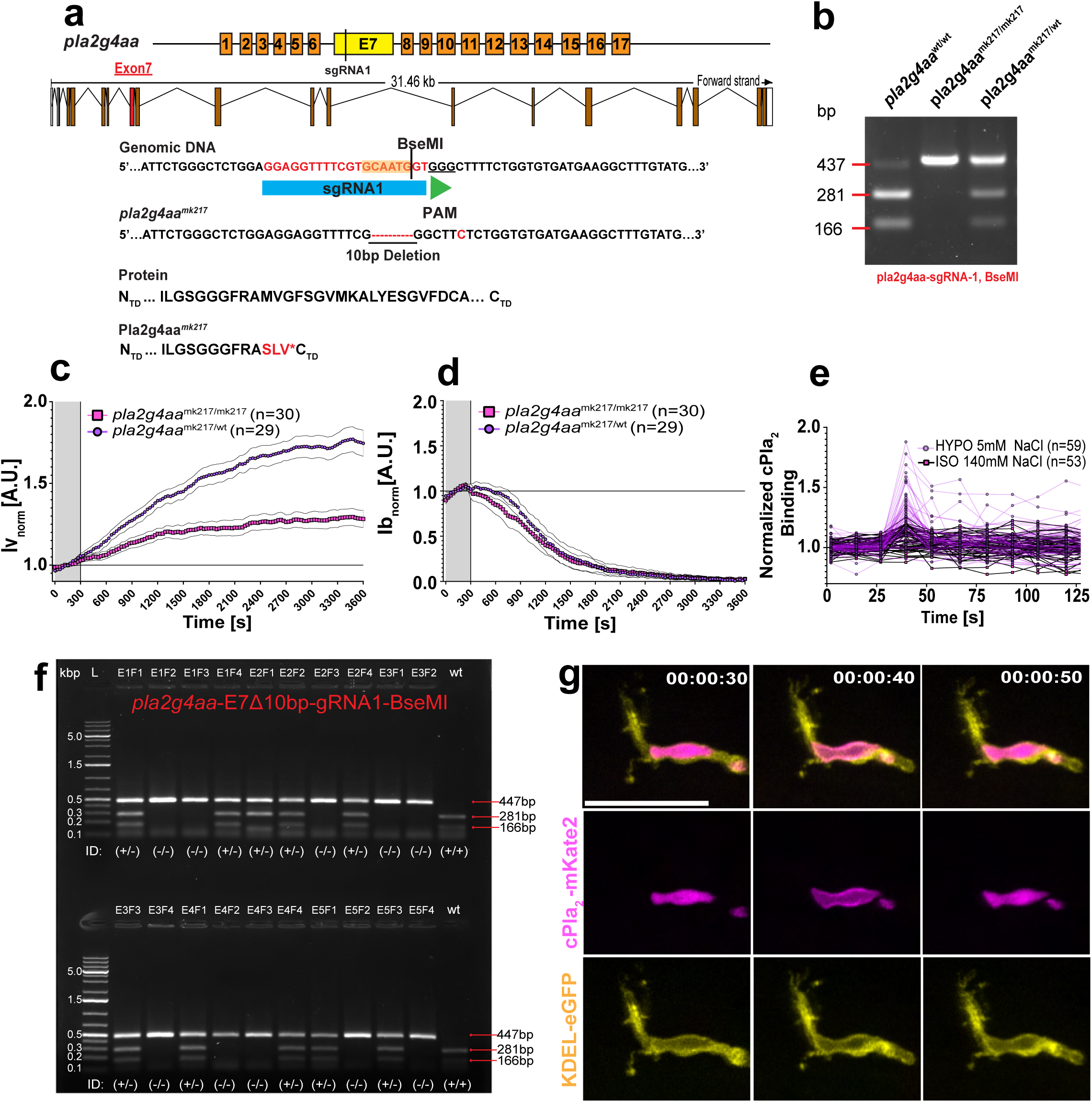
(**a**) Scheme illustrating the design of a single guide RNA targeting exon7 of *pla2g4aa* mutant zebrafish. bottom the architecture of exons (brown and red) and introns (black lines) in the zebrafish *pla2g4aa* genomic sequence. Successful gene disruption is validated with BseMI (GCAATG^GT) exon7 restriction palindrome sites (yellow box) on genomic DNA, which is altered in *pla2g4aa*^mk217/mk217^ mutants. (**b**) Representative gel genotyping of CRISPR/Cas9 induced *pla2g4aa* mutations by BseMI restriction digest. The 447 bp (PCR) product of wildtype allele is cleaved by BseMI into two smaller (281 and 166 bp) fragments. The mk217 allele carries 10bp deletion, preventing the targeted BseMI restriction enzyme digest in *pla2g4aa*^mk217/mk217^(437bp) and *pla2g4aa^mk2^*^17^*^/wt^*(437, 281 and 166bp). (**c**) Normalized vessel-(Iv_norm_(T)) and (**d**) wound-(Ib_norm_ (T)) leakage dynamics after ISO_(NaCl)_ → HYPO shifting at the indicated genetic conditions. Error margin, 95% confidence interval. (**e**) Individual plots underlying Figure 4f. (**f**) Representative genotyping for *pla2g4aa*^mk217/wt^ and *pla2g4aa*^mk217/mk217^ embryos after image acquisition. Red arrows, restriction enzyme cleavage patterns. (**g**) Representative, pseudo-coloured cPla_2_-mKate2 and endoplasmic reticulum (KDEL-eGFP) fluorescence in a perivascular macrophage before (00:00:30) and after (00:00:40 & 00:00:50) laser injury in hypotonic E3 solution. Timestamp, hh:mm:ss. Scale bars, 25 μm.

**Figure S6. Extended data supporting Fig. 4.**
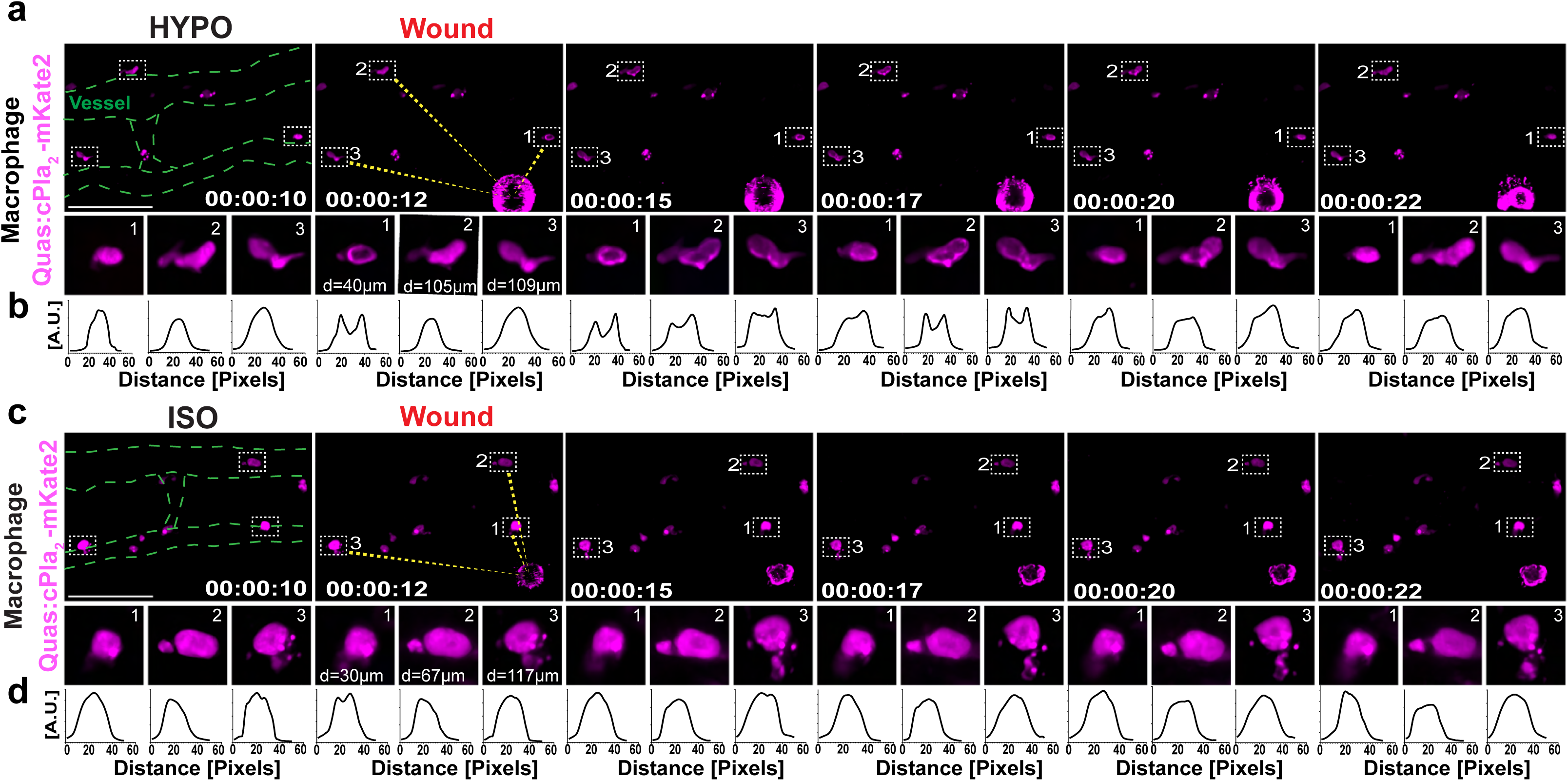
Representative two-photon montage of live Tg(*mpeg1.1*:Qf2;Quas:cPla2-mKate2) larvae imaged with rapid temporal resonant scanning (T-step=2.5 s, 6 fps) upon laser injury at T=12 s in (**a**) HYPO and (**c**) ISO_(NaCl)_ E3 bathing solution. Magenta, pseudo-coloured for cPla_2_-mKate2. Images are representative of three independent wounding experiments. Dashed green line, vessel outline. Timestamp, hh:mm:ss. Scale bars, 50 μm. (**b, d**) Line profiles for cPla_2_-mKate2 emission([A.U.]) over distance(Pixels) through the macrophages in the numbered ROIs.

## Supplementary Videos Legends

**Supplementary Video 1.** Representative time lapse movie of a 2 dpf zebrafish Tg(*kdrl*:eGFP)-larva showing vessel dilation and vessel permeability upon shifting of bathing solutions from ISO to HYPO. Overexposure of dextran emission is to highlight the bleed out from the wound. Magenta, 70 kDa dextran. Green, vessels. Timestamp, hh:mm:ss. Scale bars, 50 μm.

**Supplementary Video 2.** Representative time lapse movie of a dextran-injected and – wounded 2 dpf Tg(*kdrl*:eGFP)-larva showing inhibition of vessel dilation and permeability upon shifting of bathing solutions from ISO to ISO_NaCl_. Magenta, 70 kDa dextran. Green, vessels. Timestamp, hh:mm:ss. Scale bars, 50 μm.

**Supplementary Video 3.** Representative time lapse movie of a dextran-injected and – wounded 2 dpf Tg(*kdrl*:eGFP)-larva showing mild inhibition of vessel dilation and permeability upon shifting of the bathing solution from ISO to ISO_ChCl_. Magenta, 70 kDa dextran. Green, vessels. Timestamp, hh:mm:ss. Scale bars, 50 μm.

**Supplementary Video 4.** Representative time lapse movie of a dextran-injected and – wounded 3 dpf Tg(*lyz*:NTR2.0)-larva showing that neutrophil depletion does not impact vessel-or wound-permeability under hypotonic conditions. Green, neutrophils. Magenta, 70 kDa dextran. Timestamp, hh:mm:ss. Scale bars, 50 μm.

**Supplementary Video 5.** Representative time lapse movie of a dextran-injected and – wounded 3 dpf Tg(*mpeg1.1*:NTR2.0)-larva showing that absence of macrophages inhibits vessel leakage without impacting wound permeability. Green, macrophage. Magenta dextran 70 kDa. Timestamp, hh:mm:ss. Scale bars, 50 μm.

**Supplementary Video 6.** Representative time lapse movie of cPla_2_-mKate2 emission in macrophage nuclei under HYPO and ISO treatments, showing membrane binding after UV-laser wounding (at T= 00:00:40) and rapid recovery (at T= 00:00:50) in HYPO bathing condition. Timestamp, hh:mm:ss. Scale bars, 50 μm.

**Supplementary Video 7.** Representative two-photon imaging of macrophage cPla_2_-mKate2 emission with fast temporal acquisition in HYPO and ISO treated zebrafish larvae showing that rapid and reversible cPla_2_-INM adsorption propagates in wave-like fashion from the site of laser injury (at T=00:00:12). Timestamp, hh:mm:ss. Scale bars, 50 μm.

## Notes

### Competing Interest Statement

The authors have declared no competing interest.

